# Olfactory Mucosa-Derived Mesenchymal Stem Cells Differentiate Towards a Schwann Cell-Like Phenotype Towards Sourcing for Peripheral Nerve Regeneration

**DOI:** 10.1101/2024.09.03.611001

**Authors:** Katelyn Neuman, Abigail N. Koppes, Ryan A. Koppes

**Affiliations:** Department of Chemical Engineering, Northeastern University, Boston, Massachusetts, 02115, USA; Department of Bioengineering, Northeastern University, Boston, Massachusetts, 02115, USA; Department of Biology, Northeastern University, Boston, Massachusetts, 02115, USA

## Abstract

Mesenchymal stem cells (MSCs) are a promising source of stem cells for treating peripheral nerve injuries. Here, we present the first investigation of differentiation of olfactory mucosa-derived MSC (OM-MSC) towards a Schwann Cell (SC)-like phenotype. OM-MSCs are an advantageous potential source of SCs for peripheral nerve repair, as isolation can be accomplished with a minimally invasive procedure compared to autologous nerve harvest and isolation. Here, Schwann Cell Conditioned Media (SCCM) or a defined growth factor supplemented media (GF) was applied to OM-MSC for twenty-one days. The differentiation process and resulting populations were characterized by immunocytochemistry and RT-qPCR. Functionality of differentiated populations was assessed in an *in vitro* co-culture model to evaluate interaction with sensory neurons (dorsal root ganglia) juxtaposed to native SCs. Compared to undifferentiated MSCs, differentiation protocols resulted in significant changes in morphology, gene expression, and functionality using SCCM and GF media, representing key characteristics of SCs. Specifically, differentiated populations exhibit elongated, spindle-like morphologies, a high degree of eccentricity, increased S-100, CD44, and NGF expression, and colocalization of myelin basic proteins with neurites in the co-culture model. In conclusion, this work highlights the potential of OM-MSCs to be expanded and differentiated to SCs to improve synthetic scaffolds or for use in decellularized allografts for nerve repair.

---

Injuries to the peripheral nervous system are often caused by trauma to the extremities. Leading causes include car accidents, gunshot injuries, and stretching/crush injuries, with a high (83%) prevalence in people under the age of 55^1^. Peripheral nerve injuries (PNI) often result in loss of sensation, function, pain, and paralysis^2,3^. Despite current treatment options, many patients suffer from chronic pain and a reduced functional capacity^3-6^. Surgical interventions are needed for large gap injuries (nerve gaps greater than 3 cm). The gold standard of treatment remains to be the implantation of a nerve autograft harvested from the sural nerve to bridge the gap between the distal and proximal nerve stumps. The fundamental justification for the lack of success of engineered grafts on the market is the lack of supportive Schwann Cells (SCs). Without SCs in injury gaps greater than 1 cm, axons do not regenerate sufficiently across the gap, and patients never see a functional recovery. In large gap injuries, the native SCs (from the distal/proximal nerve stumps) become stressed or exhausted, unable to keep up with the high proliferation demand^7^. These cells then negatively affect the microenvironment, impinging regeneration by releasing inflammatory cytokines^7^. Further, autografting is undesirable, as it requires secondary surgery, damaging a secondary nerve and increasing the patient’s risk for infection. Therefore, an abundant source of a patient’s SCs is paramount to the success of new repair technologies.

Decellularized cadaver tissue allografts provide topographical cues similar to that of the native peripheral nerve extracellular matrix (ECM) and are growing in popularity for small gap injuries^8^. While removing the need for secondary surgery, these allografts have no support cells, including SCs. SCs reside in the peripheral nerve and are fundamental in forming and repairing damaged tissue, producing neurotrophic factors, and remyelinating^9^. In the presence of injury, the phenotypes of SCs alter from their myelinated state to a progenitor-like cell^10-12^. This change promotes SCs to secrete neurotrophic factors, clear damaged myelin, express adhesive and axonal cues, initiate an inflammatory response, and proliferate^10^. These characteristics support nerve repair and regeneration after an injury has occurred. While beneficial, SCs are also invasive to retrieve - with similar drawbacks of harvesting an autograft, requiring a nerve biopsy, which often results in donor site morbidity^13^. Because SCs are limited, the cells are cultured and expanded in vitro for extended periods^14^. When sufficient cells are obtained, these cells can be delivered to the injury site in combination with a scaffold material to facilitate nerve regeneration.

Schwann-like cells derived from stem cells have been investigated as an alternative. Recent research has focused on mesenchymal stem cells (MSCs) collected from different tissue sources such as adipose tissue, human dental pulp, bone marrow, olfactory neuroepithelial tissue, and skeletal muscle tissue. MSCs are attractive as support cell options for peripheral nerve repair because of their anti-inflammatory properties and ability to differentiate into neural tissue^15,16^. Mesenchymal stem cells (MSCs), both undifferentiated and differentiated, have been demonstrated to modulate the immune response in ongoing and completed clinical trials and are thought to be immune-privileged, allowing for allogeneic use^17-20^.

Though these advantages of MSCs have been reported, the original tissue location, donor age, culture conditions, and isolation method must be investigated to predict cell behavior 15. The most widely used MSCs are derived from bone marrow (BM-MSCs) and adipose tissue (AD-MSCs). BM-MSCs have been shown to support nerve regeneration^21,22^. However, the procedure for obtaining bone marrow-derived stem cells (BMSCs) is invasive and painful, and it has a low yield of stem cells^21,23^. AD-MSCs are more abundant and less invasive to retrieve, although they have not been reported to express neural/glial genes^24,25^. Therefore, BM-MSCs and AD-MSCs may not be ideal for PNIs.

In this work, we utilized olfactory mucosa-derived mesenchymal stem cells (OM-MSCs). Due to their unique embryology (derived from neural crest vs. mesoderm) and niche (or original tissue location), we hypothesize that OM-MSCs have a higher potential of differentiation towards an SC-like phenotype and an increased tendency to promote peripheral nerve regeneration^26-28^. These cells have a high proliferation rate and are easily accessible from human patients with a minimally invasive outpatient procedure^29-31^. This study aims to investigate, for the first time, the differentiation of OM-MSCs towards an SC-like cell using Schwann Cell Conditioned Media (SCCM) and a growth factor cocktail (GF). This study uses immunocytochemistry, flow cytometry, and RT-qPCR to characterize the differentiation process and resulting populations. The ability to support neurite extension and exhibit the first steps towards myelination in vitro highlight the utility of OM-MSCs in driving an SC phenotype for the treatment of PNI.

## Results

### Immunocytochemistry Characterization

ICC characterization of freshly isolated OM-MSC exhibits an expected flat, fibroblast-like morphology typical of mesenchymal stem cells. OM-MSCs have robust expressions of S100β, cluster of differentiation 105 (CD105), and actin filaments (Figure 1). Most OM-MSCs express Nestin, and all cells appear to have low-level expression of βIII tubulin and GFAP (Supplementary Fig. 1). SCs (Figure 3) and astrocyte (Supplementary Fig. 2) populations for the following experiments were determined to be homogenous.

**Fig. 1.**
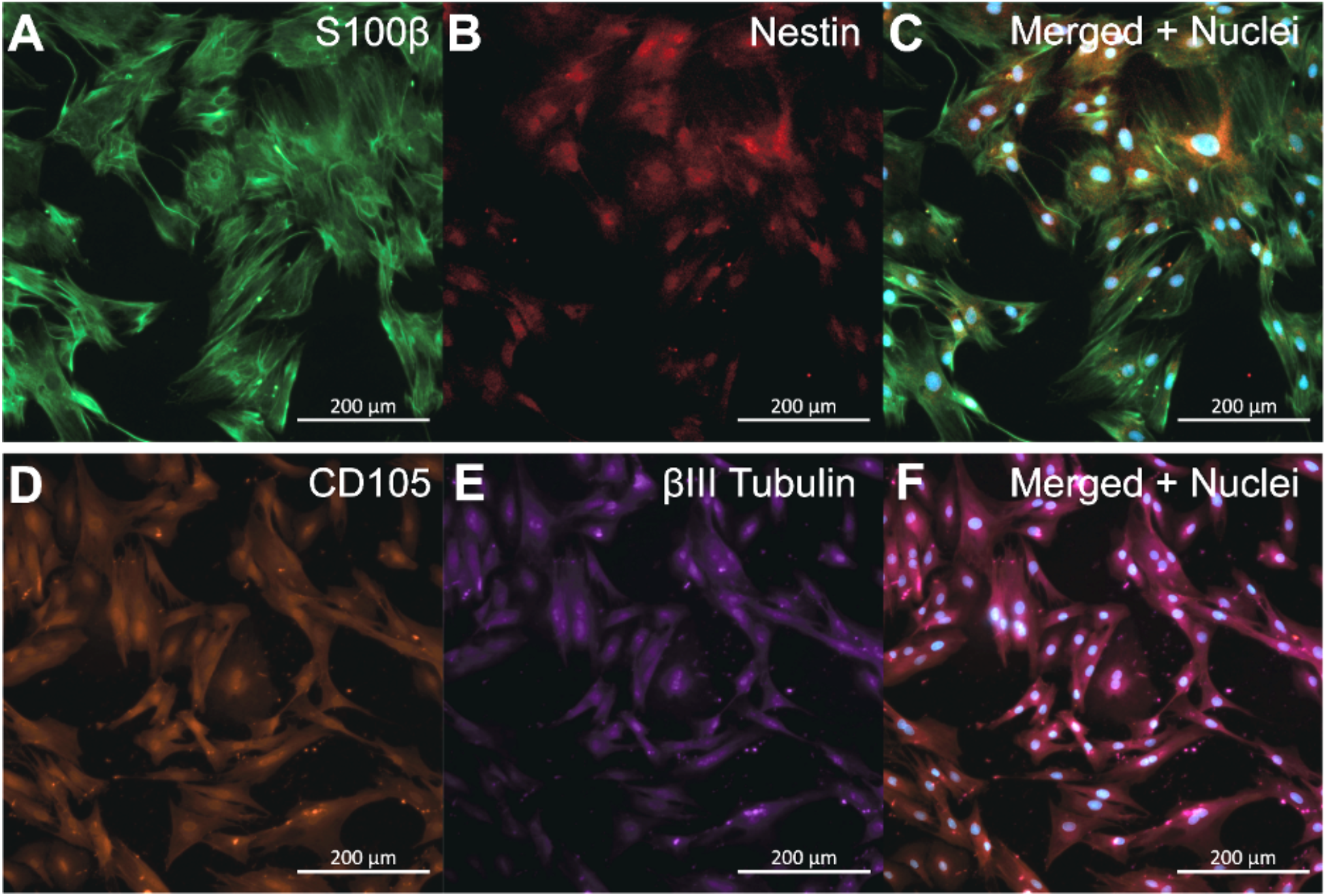
Characteristic immunocytochemistry of OM-MSCs, isolated from rat tissue, display flat, fibroblast-like morphology typical of mesenchymal cells. (A) Green fluorescence denotes S100β, (B) Red fluorescence denotes Nestin, and (C) Displays both S100β and Nestin expression merged with cell nuclei (shown in blue, counterstained with DAPI). (D) Orange fluorescence denotes CD105, (E)Purple fluorescence denotes βIII tubulin, and (F) Displays both CD105 and βIII Tubulin expression merged with cell nuclei (shown in blue, counterstained with DAPI). Scale Bar = 200 µm.

### OM-MSCs Express Mesenchymal Stem Cell Markers

OM-MSCs isolated from female adult Sprague Dawley rats were assessed in a Beckman Coulter CytoFLEX for markers outlined in Table 2. Cell populations were selected by removing debris, doublets, and dead cells before analysis (Figure 2A-C). Less than 1% of OM-MSCs expressed cluster of differentiation 45 (CD45), a hemopoietic stem cell marker and negative control (Figure 2D). 99.8% and 94% of OM-MSCs expressed mesenchymal stem cell markers CD44 and CD90, respectively (Figure 2E-F). 60% of OM-MSCs expressed Nestin, a marker of stemness (Figure 2G). All cells that expressed Nestin were also CD44^+^ (Figure 2H). <3% of cells that were Nestin^+^, did not express CD90 (Figure 2J). Similar results were recorded for OM-MSCs isolated from male Sprague Dawley rats (Supplementary Fig. 3).

**Fig. 2.**
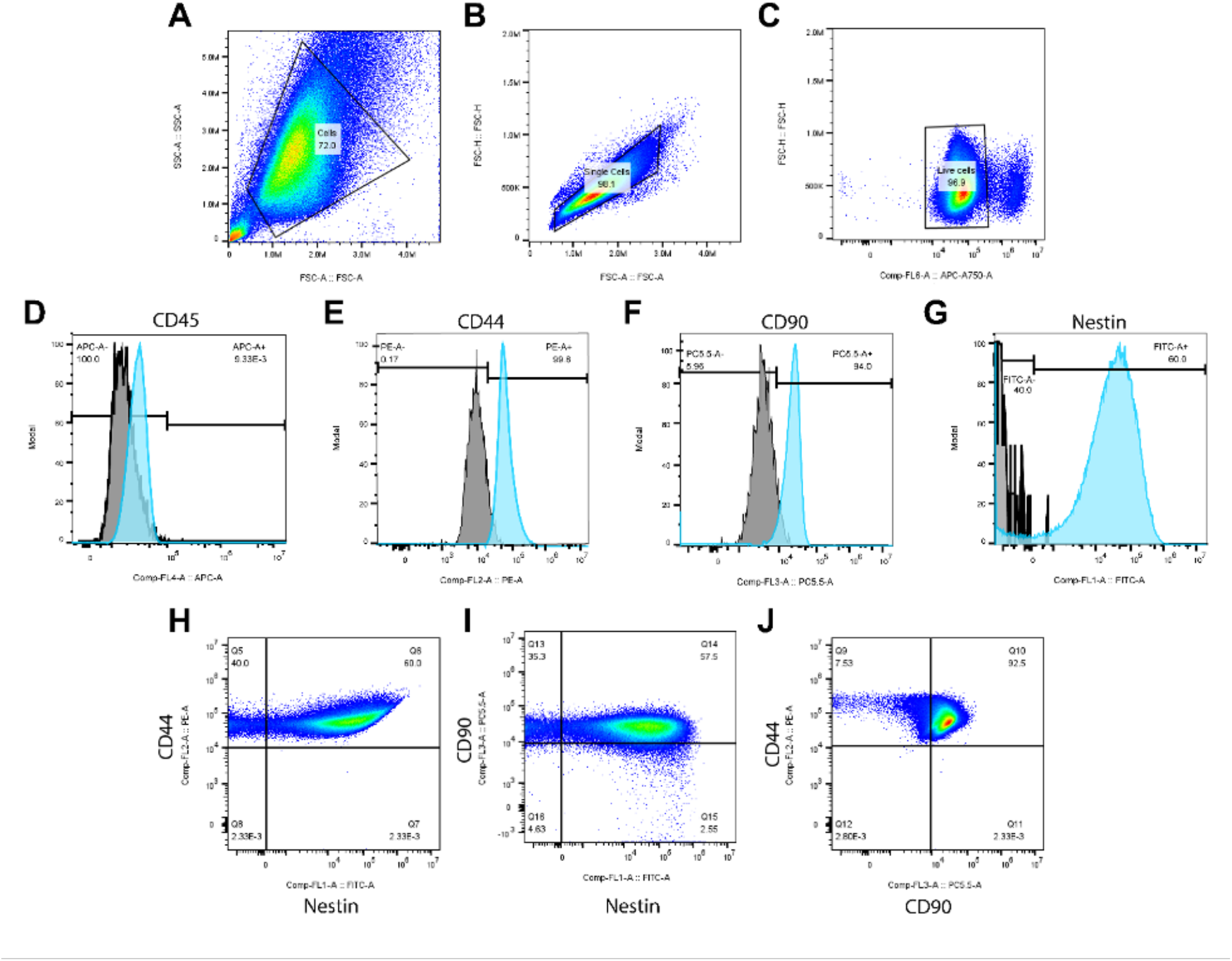
Characteristic flow cytometry of OM-MSCs. (A-C) Display gating strategy employed prior to analysis. The selection of (A) cell population and removal of debris, (B) single cells, and then (C) live cell population is shown. (D-G) Display the distribution of positive and negative cells for the four markers of interest, (D) CD45, (E) CD44, (F) CD90, and (G) nestin. The gray histogram represents the negative control, and the blue histogram represents the cells of interest. (H-I) Display the co-localization of (H) CD44 and nestin expression, (I) CD90 and nestin expression, and (J) CD44 and CD90 expression.

**Fig. 3.**
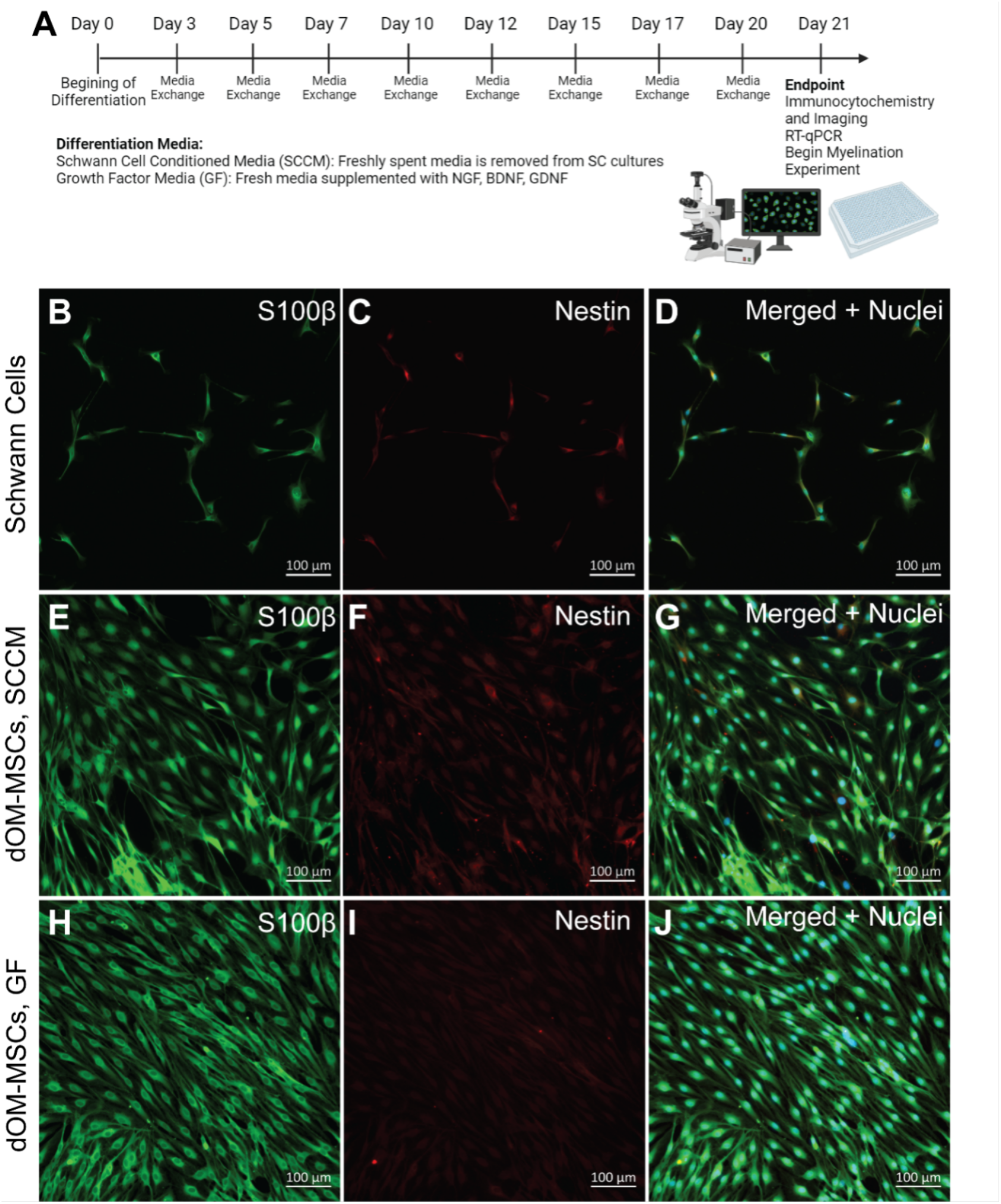
Differentiated OM-MSC morphology resembles the small, spindle shape of SCs. (A) 21-day differentiation timeline of OM-MSCs. Created with Biorender.com (B-D) Representative images of SCs with (B) denoting S100β (green fluorescence), (C) denoting nestin (red fluorescence), and (D) displaying both S100β and nestin expression merged with cell nuclei (shown in blue, counterstained with DAPI). (E-G) Representative images of dOM-MSCs with SCCM with (E) denoting S100β (green fluorescence), (F) denoting nestin (red fluorescence), and (G) displaying both S100β and nestin expression merged with cell nuclei (shown in blue, counterstained with DAPI). (H-J) Representative images of dOM-MSCs with GF media with (H) denoting S100β (green fluorescence), (I) denoting nestin (red fluorescence), and (J) displaying both S100β and nestin expression merged with cell nuclei (shown in blue, counterstained with DAPI). Scale bar = 100 µm.

### Differentiated OM-MSCs Resemble SCs

Cultured for 21 days in both SCCM and GF media, dOM-MSCs exhibited cell morphology and Nestin expression changes. The dOM-MSCs retained S100β expression (Figure 2). Other media formulations were then evaluated to optimize the differentiation process. OM-MSCs did not differentiate when subjected to a previously published chemical induction protocol applied to BM-MSCs 39. Also, less forskolin in the growth factor media resulted in a reduction in morphology changes and Nestin expression at day 14 (Supplementary Fig. 4). Cell Profiler™ was used to quantify these changes in cell morphology and compare them directly to undifferentiated OM-MSCs and SCs. The cell area of SCs, OM-MSCs, dOM-MSCs (SCCM), and dOM-MSCs (GF) were measured to be 490 ± 270 *µ*m^2^, 3900 ± 2500 *µ*m^2^, 400 ± 210 *µ*m^2^, and 1000 ± 410 *µ*m^2^ respectively (Figure 4A). All groups were significantly different from each other with all comparisons, having a p-value < 0.0001, except SC vs. dOM-MSCs (SCCM, p = 0.009). The eccentricity was also measured. SCs, OM-MSCs, dOM-MSCs (SCCM), and dOM-MSCs (GF) were estimated to be 0.90 ± 0.11, 0.76 ± 0.14, 0.81 ± 0.16, and 0.88 ± 0.12, respectively (Figure 4B). Again, all groups were significantly different, with a p-value < 0.0001, apart from SC vs dOM-MSCs (GF, p = 0.003). The median radius of each cell body of SCs, OM-MSCs, dOM-MSCs (SCCM), and dOM-MSCs (GF) measured to be 2.1 ± 0.6 *µ*m, 5.4 ± 2.2 *µ*m, 2.0 ± 0.5 *µ*m, and 2.8 ± 0.6 *µ*m, respectively (Figure 4C). SCs and dOM-MSCs (SCCM) were not significantly different from each other (p >0.9999). All others were significantly different from each other (p < 0.0001). Finally, the cell perimeter was also measured with SCs, OM-MSCs, dOM-MSCs (SCCM), and dOM-MSCs (GF) having perimeters of 160 ± 87 *µ*m, 710 ± 410 *µ*m, 140 ± 79 *µ*m, and 240 ± 99 *µ*m, respectively (Figure 4D). SCs and dOM-MSCs (SCCM) were not significantly different (p = 0.32). All others were significantly different from each other (p < 0.0001).

**Fig. 4.**
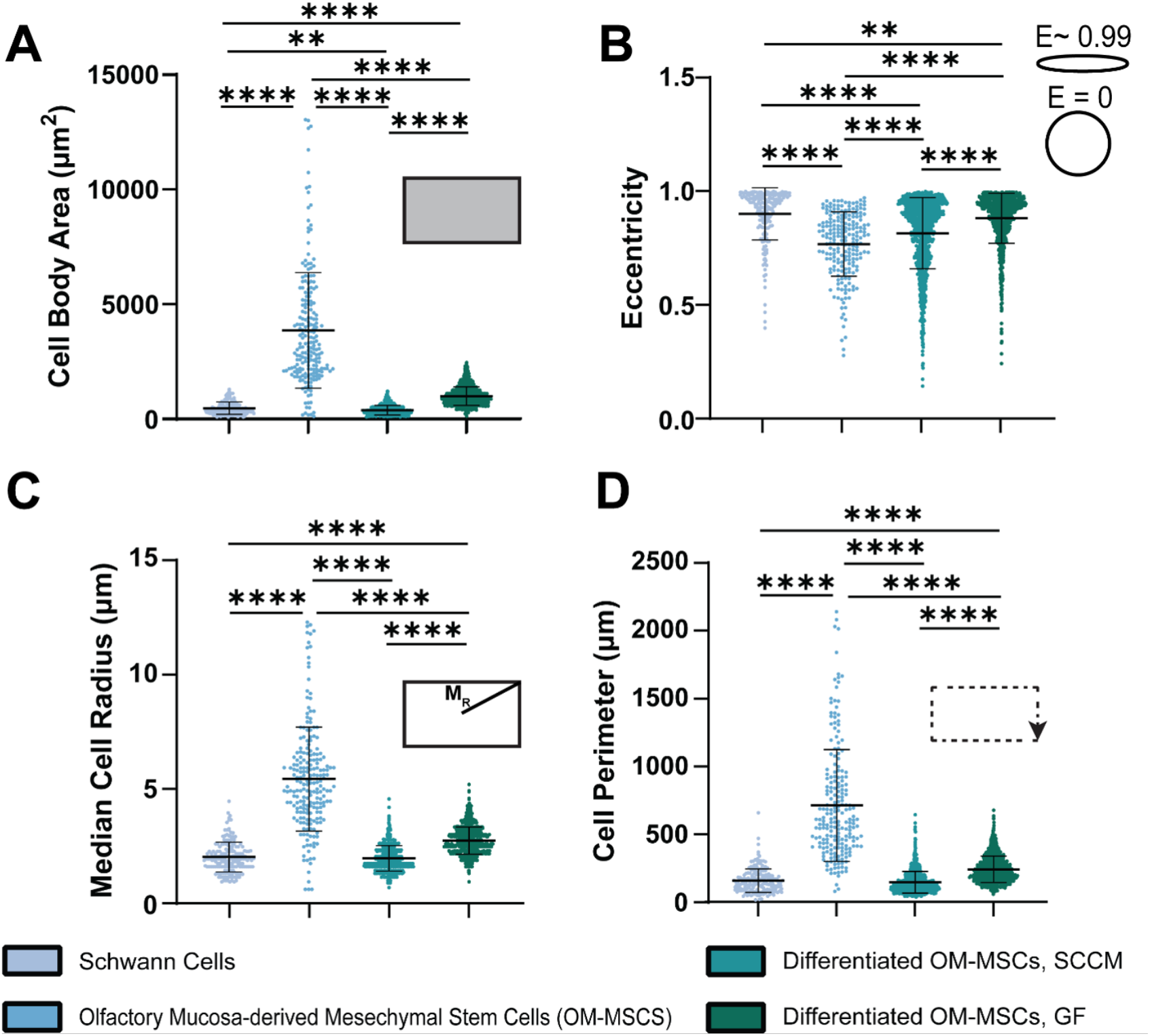
Changes in cell shape in differentiated OM-MSCs (dOM-MSCs) for both SCCM and GF media conditions were compared to SCs and OM-MSCs using Cell Profiler40. Results for (A) cell area, (B) eccentricity, (C) median cell radius, and (D) cell perimeter. Ten images were analyzed for each condition (n = 190, 210, 1434, 1621). A Krustal-Wallis test (non-parametric ANOVA) with Dunn’s multiple comparisons test was used to assess significance. ** = p< 0.01 and **** = p<0.0001. Error bars display the standard deviation.

### Gene expression of dOM-MSCs in SCCM resembles that of SCs

Relative to undifferentiated OM-MSCs, the degree of expression of ten different genes across five experimental groups was evaluated with RT-qPCR (Figure 5). Relative expression of CD44 was upregulated in DRG (log2fold change of 4.5 ± 3.1) compared to OM-MSCs (log2fold change of 0.0 ± 0.5, p = 0.03) and dOM-MSCs (GF, log2fold change of -1.2 ± 2.5, 0.005, Figure 5A). No significant differences in relative gene expression of CD90 across all groups were measured (Figure 5B). The relative expression of GDNF was significantly downregulated in astrocytes (log2fold change of -13.4 ± 4.3) compared to all other groups (p < 0.0001, Figure 5C). GFAP was upregulated in astrocytes (log2fold change of 13.1 ± 0.8), which was significantly more than DRG (log2fold change of 4.2 ± 2.3, p = 0.0003), OM-MSCs (log2fold change of 0.0 ± 2.1, p < 0.0001), dOM-MSCs (GF, log2fold change of 6.0 ± 4.2, p = 0.004), and dOM-MSCs (SCCM, log2fold change of -6.8 ± 1.5, p < 0.0001). No significant difference in relative gene expression in GFAP was observed between astrocytes and SCs (log2fold change of 9.0 ± 0.7, p = 0.16).

**Fig. 5.**
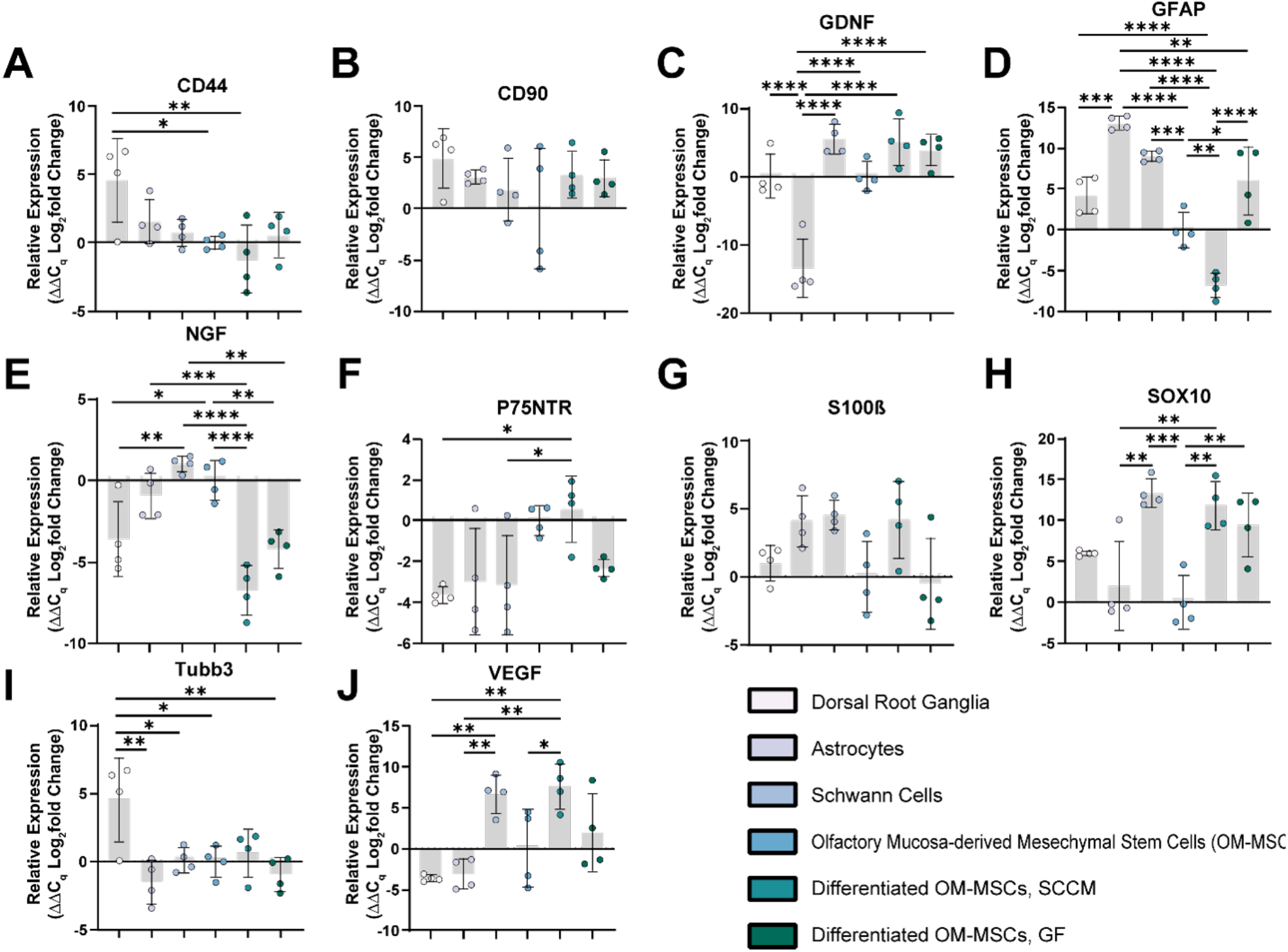
Relative gene expression of dOM-MSCs in SCCM closely resembles that of SCs. Relative log2fold changes in gene expression of (A) CD44, (B) CD90, (C) GDNF, (D) GFAP, (E) NGF, (F) P75NTR, (G) S100β, (H) SOX10, (I) Tubb3, and (J) VEGF. Gene expression relative to Olfactory Mucosa-derived Mesenchymal Stem Cells (OM-MSCs) was compared across five experimental groups: Dorsal Root Ganglia, Astrocytes, Schwann Cells (SCs), differentiated OM-MSCs (SCCM), and differentiated OM-MSCs (GF). Four independent samples were assessed per condition. Significance was assessed with a one-way ANOVA and Tukey’s multiple comparison test. Significance is denoted with *, **, ***, and **** which represent p values of < 0.05, <0.01, <0.001, and <0.0001. Error bars indicate standard deviation.

Compared to OM-MSCs, relative GFAP expression was upregulated in SCs (p = 0.0003) and dOM-MSCs (GF, p = 0.02). Relative GFAP expression was significantly downregulated in dOM-MSCs (SCCM) compared to DRG (p < 0.0001), SCs (p < 0.0001), OM-MSCs (p = 0.006), and dOM-MSCs (GF, p < 0.0001, Figure 5D). Relative expression of NGF was observed to be downregulated in DRG (log2fold change of -3.6 ± 2.3, p = 0.03), dOM-MSCs (SCCM, log2fold change of -6.8 ± 1.5, p < 0.0001), dOM-MSCs (GF, log2fold change of -4.2 ± 1.2, p = 0.01) compared to OM-MSCs (log2fold change of 0.0 ± 1.2). When compared to SCs, the relative expression of NGF was significantly downregulated in DRG (p = 0.003), dOM-MSCs (SCCM, p < 0.0001), and dOM-MSCs (GF, p = 0.001). NGF was significantly upregulated in astrocytes (log2fold change of -0.9 ± 1.4) compared to dOM-MSCs (SCCM, p = 0.0003, Figure 5E).

P75NTR was upregulated in dOM-MSCs (log2fold change of 0.7 ± 1.6) compared to DRG (log2fold change of -3.6 ± 0.4, p = 0.02) and SCs (log2fold change of -3.1 ± 2.4, p = 0.05, Figure 5F). No significant differences in relative gene expression of S100β were observed, although SCs and dOM-MSCs (SCCM) had similar levels of relative expression, with log2fold changes of 4.5 ± 1.1 and 4.2 ± 2.8 respectively (p > 0.9999, Figure 5G). SOX10 was upregulated in SCs (log2fold change of 13.3 ± 1.7), dOM-MSCs (SCCM, log2fold change of 11.8 ± 2.9), and dOM-MSCs (GF, log2fold change of 9.4 ± 3.9) compared to OM-MSCs (log2fold change of 0.0 ± 3.2, p = 0.0003, 0.001, 0.008). The upregulation of SOX10 expression observed in SCs and dOM-MSCs (SCCM) was also significantly different than the relative expression level in astrocytes (log2fold change of 2.0 ± 5.4, p = 0.002, 0.006, Figure 5H). DRG were observed to have a significant upregulation in the relative expression of Tubb3 (log2fold change of 4.5 ± 3.1) compared to astrocytes (log2fold change of -1.5 ±1.6, p = 0.002), SCs (log2fold change of 0.1 ± 0.8, p = 0.02), OM-MSCs (log2fold change of 0.0 ± 1.1, p = 0.02), and dOM-MSCs (GF, log2fold change of -1.0 ± 1.2, p = 0.004). A trending up regulation of Tubb3 expression was also observed in DRG compared to dOM-MSCs (SCCM, log2fold change of 0.6 ± 1.7, p = 0.05, Figure 5I). VEGF was significantly upregulated in SCs (log2fold change of 6.7 ± 2.3) when compared to the relative expression levels in astrocytes (log2fold change of -3.1 ± 1.9, p = 0.005) and DRG (log2fold change of -3.6 ± 0.4, p = 0.003). VEGF was also significantly upregulated in dOM-MSCs (SCCM, log2fold change of 7.6 ± 2.7) when compared to astrocytes (p = 0.002), DRG (p = 0.001), and OM-MSCs (log2fold change of 0 ± 4.7, p = 0.04, Figure 5J).

### Differentiated OM-MSCs exhibit Co-Localized Expression of MBP with extending Neurites from dissociated sensory neurons

The ability of the differentiated cells to perform a crucial function of SCs and myelinate DRG was assessed (Figure 6). Representative images of each of the four groups assessed, SCs, OM-MSCs, dOM-MSCs (SCCM), and dOM-MSCs (GF) are displayed in Figure 6B-M. The percent of MBP overlapping NF-H of each was measured to be 67 ± 17%, 47 ± 23%, 64 ± 19%, and 65 ± 20%, respectively. The groups with SCs, dOM-MSCs (SCCM), and dOM-MSCs (GF) as support cells were all significantly different from those with OM-MSCs (p = 0.0002, 0.003, 0.0006). Those three groups had no significant differences (p > 0.9999, Figure 6N). The Mander’s coefficient of each group was also measured and found to be 0.68 ± 0.17, 0.48 ± 0.23, 0.64 ±.19, and 0.65 ± 0.20, respectively. Like the previous analysis, the groups with SCs, dOM-MSCs (SCCM), and dOM-MSCs (GF) as support cells were all significantly different from those with OM-MSCs (p = 0.0001, 0.004, 0.0008). There were no significant differences between SCs, dOM-MSCs (SCCM), and dOM-MSCs (GF, p >0.9999, Figure 6O).

**Fig. 6.**
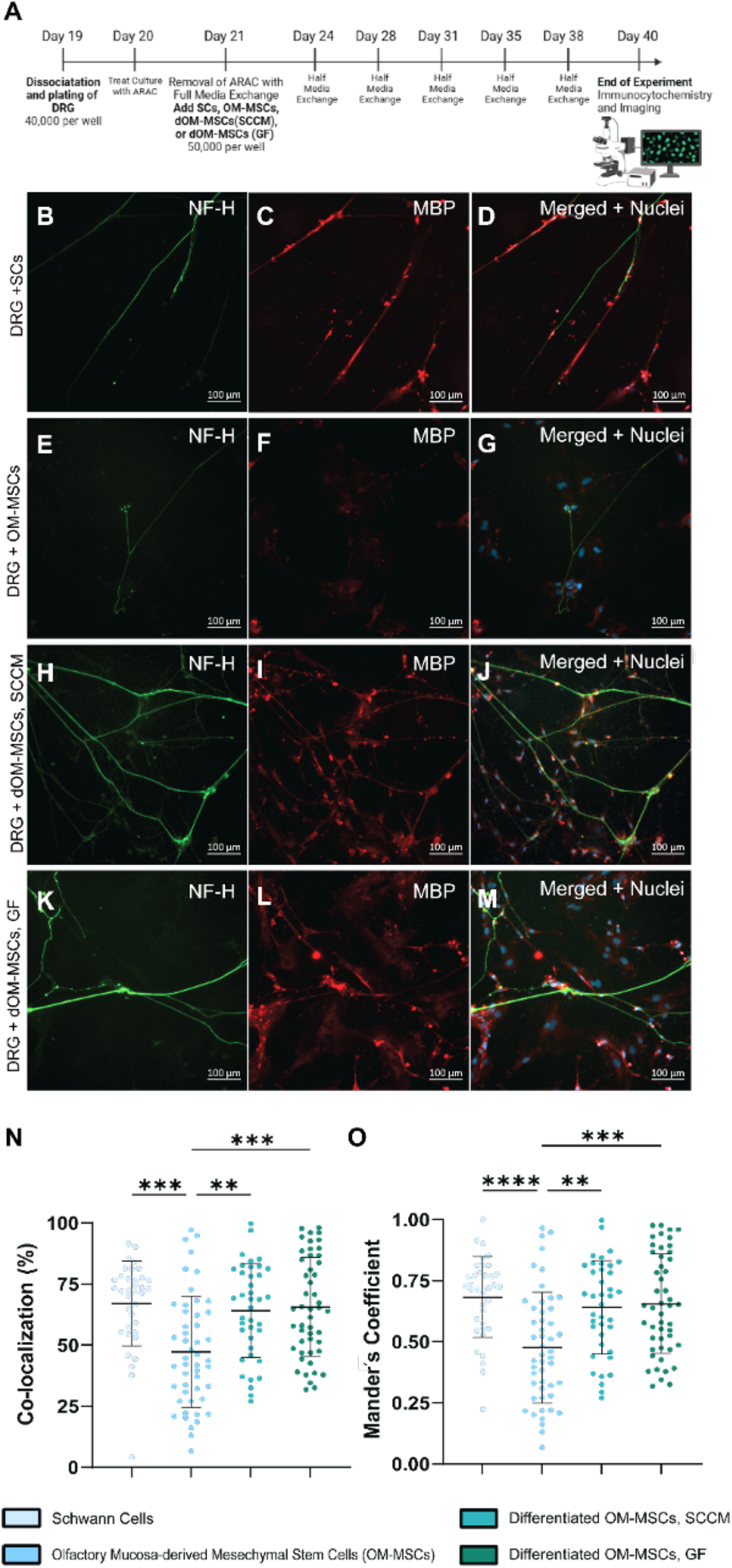
Differentiated OM-MSCs exhibit co-localized expression of MBP on DRG neurons comparable to SCs. (A) 21-day experimental timeline. Created with Biorender.com. (B-D) Representative images of SCs with dissociated DRG with (B) denoting neurofilament-heavy (NF-H; green fluorescence), (C) denoting myelin basic protein (MBP; red fluorescence), and (D) displaying both NF-H and MBP expression merged with cell nuclei (shown in blue, counterstained with DAPI). (E-G) Representative images of OM-MSCs with dissociated DRG with (E) denoting NF-H (green fluorescence), (F) denoting MBP (red fluorescence), and (G) displaying both NF-H and MBP expression merged with cell nuclei (shown in blue, counterstained with DAPI). (H-J) Representative images of dOM-MSCs (SCCM) with dissociated DRG with (H) denoting NF-H (green fluorescence), (I) denoting MBP (red fluorescence), and (J) displaying both NF-H and MBP expression merged with cell nuclei (shown in blue, counterstained with DAPI). (K-M) Representative images of dOM-MSCs (GF) with dissociated DRG with (K) denoting NF-H (green fluorescence), (L) denoting MBP (red fluorescence), and (M) displaying both NF-H and MBP expression merged with cell nuclei (shown in blue, counterstained with DAPI). Scale bar = 100 µm. (N) Displays the percent of neurites that had co-localized expression of MBP. (O) Displays the Mander’s coefficient for each of the four experimental groups. Three independent trials were conducted with at least 36 images analyzed for each condition (n = 36, 49, 29, 49). A Krustal-Wallis test (non-parametric ANOVA) with Dunn’s multiple comparisons test was used to assess significance. ** = p< 0.01 and *** = p<0.001. Error bars display the standard deviation.

## Discussion

The olfactory system, which is responsible for our sense of smell, has uniquely evolved to continually support nerve regeneration due to its constant exposure to the outside environment^46-49^. This constant neurogenesis is fueled by stem cells in the surrounding tissue50. Multipotent stem cells that reside in the lamina propria of the olfactory mucosa have been identified previously as OM-MSCs due to their predicted neuroectoderm embryonic origin^31,50-53^. Isolating the lamina propria of the olfactory mucosa yielded plastic-adherent cells with fibroblast-like morphology (Figure 1, Supplementary Fig. 1), typical of MSCs from various sources and species^54-57^. OM-MSCs exhibited expression of three known MSC stem cell markers, CD44, CD90, and CD105, but not a hematopoietic stem cell marker, CD45 (Figure 1, Figure 2)^51,54,56,58-61^. CD44 is a cell surface receptor for glycosaminoglycan hyaluronan that mediates cell adhesion to the extracellular matrix^62^. CD90 (Thy-1) is a cell membrane glycoprotein that may play a role in MSCs self-renewal and differentiation^63,64^. CD105 (endoglin) is another cell membrane protein component of the receptor complex of transforming growth factor-beta. This protein is involved in cell proliferation, differentiation, and migration^65^. Lastly, CD45 is a cell surface marker of tyrosine phosphatase, an exclusive marker of hematopoietic lineage cells66. Here, our isolated OM-MSCs were confirmed to be characteristically MSCs.

In addition to the traditional MSCs, we identified the expression of several neural and glial cell-specific proteins expressed by the OM-MSCs before differentiation. There was a low-level expression of βIII tubulin (also known as Tuj-1), a standard neuronal marker, a microtubule found in the cytoskeletal of developing and regenerating axons in undifferentiated OM-MSCs (Figure 1)^67^. GFAP was also minimally expressed in OM-MSCs (Supplementary Fig. 1). GFAP is a known marker for mature astrocytes, one type of central nervous system glial cell68. Strong S100β and P75NTR expression was observed in the undifferentiated OM-MSCs, two common SC markers (Figure 1, Figure 5F)^69,70^. S100β and other S100 proteins involved in calcium binding regulate various cellular functions71. In the peripheral nervous system, P75NTR is an NGF receptor crucial for regulating axonal growth and myelination, among its other functions^72,73^. Using flow cytometry and immunocytochemistry, we also observed that most OM-MSCs are made of Nestin^+^ (Figure 1, Figure 2). As a neuroepithelial stem cell protein, Nestin is a cytoskeletal intermediate filament thought to play a role in stemness and differentiation, particularly for neural crest-derived stem cells^74,75^. Previous work has also demonstrated that OM-MSCs express a wide variety of neuronal/glia markers such as S100A4, GFAP, P75NTR, MAP2, βIII tubulin, Nestin, NGF, and GDNF^51,55,58-60,76^. Notably, our population of Nestin^+^ cells was substantially lower than that previously identified by Lindsay et al. (100%) and Delorme et al. (100%)^29,31^. The decrease in expression may be due to differences in cell culture conditions. Specifically, serum in cell media inhibits Nestin expression or the use of earlier passage cells^77,78^. The expression of these neural-lineage-specific genes highlight the potential for OM-MSCs to be an ideal source of SC-like cells to improve peripheral nerve repair.

Undifferentiated OM-MSCs have been demonstrated to have neurodegenerative properties. Conditioned media and OM-MSCs enhanced the myelination of motor neurons in vitro; BM-MSCs, on the other hand, did not^79^. A clinical trial that transplanted whole olfactory mucosa biopsies (containing OM-MSCs and olfactory ensheathing cells) to lesion sites in the spinal cord observed recovering sensory and motor function^80^. We sought to differentiate OM-MSCs toward an SC-like cell to improve injury response. The multipotency of OM-MSCs has been demonstrated previously by inducing differentiation towards non-neuronal lineages such as liver, heart, skeletal muscle, bone, and adipose tissue.

OM-MSCs have also been demonstrated to differentiate towards neurons and central nervous system glia, astrocytes, and oligodendrocytes^50,81^. To our knowledge, only one other study investigated differentiating OM-MSCs towards an SCs-like cell using a fibrin matrix to differentiate them, which was defined by an increase in MBP expression in vitro^76^. We aimed to improve upon this by utilizing defined media components to differentiate these cells and assess their ability to function as SCs with a co-culture of sensory neurons.

Recent work has shown that conditioned media from SCs can, in varying conditions, modify angiogenesis, modify cell survival, increase neuritogenesis, and induce stem cell differentiation^82-85^. SCCM was able to alter the morphology, gene expression, and functionality of OM-MSCs to a population that can better support nerve regeneration. These results were promising but still require the harvest and expansion of SCs, which, as mentioned above, requires a nerve harvest to retrieve. So, we sought to develop a defined GF media cocktail that could mimic the resulting differentiation of the SCCM by supplementing SC media with three well-known growth factors, NGF, BDNF, and GDNF, that are widely known to be secreted by SCs37. Additionally, we supplemented the media with forskolin as it can increase cAMP levels^38^. This method also induced SC-like cell morphology, gene expression, and functionality compared to OM-MSCs.

Changes in the relative expression of various genes were used to characterize the dOM-MSCs further. In addition to SCs and OM-MSCs, DRG and astrocyte samples were utilized as controls. In whole DRG samples containing neuronal, glial, and additional stromal cells, βIII tubulin was upregulated compared to the other experimental groups (Figure 5I). As expected, relative gene expression of GFAP in the astrocyte population, the glial/support cells of the central nervous system, was upregulated compared to all other experimental populations (Figure 5D). We also see the expected relative expression of CD90 and CD44 (Figure 5A-B).

The relative expression of two SC markers, S100β and SOX10, were evaluated with RT-qPCR. No significant changes in the relative expression of S100β, a standard SC marker preferentially expressed in myelinating SCs, were measured across the various experimental populations^69^. SCs and dOM-MSCs (SCCM) were observed to have similar log2fold changes in relative expression (Figure 5G)^69^. SOX10 is a transcription factor that is a reliable marker of SCs in vitro. It is the only known marker expressed throughout the SC development process, with immature SCs, precursor SCs, non-myelinating SCs, and myelinating SCs all expressing SOX1086. Compared to undifferentiated OM-MSCs, both differentiated cell populations significantly upregulated SOX10, similar to SCs (Figure 5H). The co-upregulation in the relative expression of both S100β and SOX10 suggests that dOM-MSCs (SCCM) are induced towards myelinating SC-like cells.

Interestingly, no changes in relative gene expression of P75NTR, which is generally expressed by non-myelinating SCs and immature SCs, was observed between the various stem cell populations (OM-MSCs, dOM-MSCs [SCCM], dOM-MSCs [GF], Figure 5F)^87^. P75NTR has recently been investigated as an MSC marker, with recent works demonstrating that the subset of MSCs that are P75NTR^+^proliferate faster and have a more significant differentiation potential88-90. Between 2-40% of the MSC population was reported to be positive for P75NTR^+^ in BM-MSCs and AD-MSCs. Future work will evaluate the expression of this marker in OM-MSCs as it may play a role in the enhanced differentiation potential^91^.

NGF and GDNF are important neurotrophic factors that play a crucial role in the survival, growth, and differentiation of neurons in the peripheral nervous system^92-95^. VEGF is an important growth factor that guides re-vascularization and blood vessel formation, which is crucial after injury^96^. The expression of important SC markers and growth factors in the dOM-MSCs, particularly (SCCM), makes these cells a promising candidate for supporting peripheral nerve regeneration. In an uninjured peripheral nerve, SCs are responsible for the myelin sheath, a fatty protein facilitating signal propagation along the nerve axon^97,98^. Thus, the ability to produce MBP is a crucial indicator of cells that can function similarly to SCs. Here, this functionality was quantified with two metrics, percent co-localization, and the Mander’s coefficient, with co-localization measuring the overlap of MBP only on the neurites and the Mander’s coefficient measuring the overall overlap of both the neurites and MBP. Both dOM-MSCs (SCCM and GF) expressed MBP and co-localized with neurite extensions significantly more than undifferentiated OM-MSCs but not significantly different than SCs (Figure 6). Additionally, the expression of MBP further demonstrates the differentiation towards myelinating SC-like cells^86^. Our in vitro co-culture experiment illustrates the promise and potential for these cells to function as SCs themselves do.

While both dOM-MSCs (SCCM and GF) had comparable co-localization of myelin basic protein on neurites, cell morphology, and gene expression differences suggest potential for improving the GF cocktail. In contrast, the GF cocktail did not impact the relative expression of SC marker S100β compared to OM-MSCs. The GF cocktail significantly upregulates SOX10, a second SC marker, compared to undifferentiated OM-MSCs. The dOM-MSCs (GF) also expressed MBP after co-culture with DRG. The difference in cell morphology and gene expression is likely due to unidentified SCCM components contributing to more complete differentiation. For example, BM-MSCs that differentiated in neurons with SCCM were found to be induced by microRNA pathways^83^.

Additionally, other molecules produced by SCs may be inducing differentiation. Future work will further optimize our growth factor media to include other growth factors produced by SCs, such as VEGF, which we see is upregulated in SCs via RT-qPCR analysis (Figure 5J). Further optimization is vital to eliminate the need for SCs to differentiate OM-MSCs and develop a protocol conducive to scalability. A defined media is paramount to translating these cells to the clinic, allowing a xeno-free and scalable process.

In conclusion, we differentiated OM-MSCs towards SC-like cells and characterized their function and phenotype. Our in vitro co-culture experiment illustrates the promise and potential for these cells to function as SCs themselves do. Future work will assess the functionality of these cells in vivo and work towards translating this protocol towards differentiating human-derived OM-MSCs for enhancing peripheral nerve regeneration in patients.

## Materials and Methods

### Animal Statement

All animal work was performed with the approval of Northeastern University’s Institutional Animal Care and Use Committee (NU-IACUC; Protocol #20-0207R) and followed the NIH guide for animal use.

### Primary Cell Isolations and Immunocytochemistry Characterization

Olfactory Mucosa Derived Mesenchymal Stem Cells (OM-MSCs) were isolated from explants removed from the nasal cavities of 6-week-old Sprague Dawley rats and used to collect multipotent OM-MSCs according to a previously established protocol 32. Briefly, after euthanasia, facial muscle, and excess tissue were removed, exposing the nasal cavity. Once exposed, the olfactory turbinates were gently lifted to uncover the olfactory mucosa between the ceiling of the nasal cavity, the arc of the perpendicular plate, and the cribriform plate. Using a 26-gauge needle and forceps, the mucosa was removed and transferred into a petri dish filled with Dulbecco’s Modified Eagle Medium/Nutrient Mixture F-12 (DMEM/F-12; Invitrogen). A similar protocol has been established for humans in which a small biopsy can be taken from the patient’s nose in a quick outpatient procedure^32^.

The rat tissue was digested in dispase II solution (2.4 IU/mL; Sigma-Aldrich) for one hour at 37°C, then transferred to a collagenase I (4 mg/ mL; Sigma-Aldrich) solution for ten minutes. The tissue was then dissociated with a pipette, and the reaction was terminated with phosphate-buffered saline (PBS; Sigma-Aldrich). The samples were then centrifuged at 300 g for five minutes. After removing the supernatant, the cell pellet was resuspended in DMEM/F-12 supplemented with 10% (v/v) fetal bovine serum (FBS; Corning) and 50 U/mL -penicillin/streptomycin (P/S; Sigma-Aldrich) and plated in a 12-well plate (Thermo Fisher). Cell medium (1 mL) was added to each well and exchanged every 2-3 days. The cells were incubated in standard cell culture conditions throughout all experiments (37°C, 5% CO2). After cells reached 70-90% confluency (about 1-2 weeks), the OM-MSCs were passaged according to standard cell culture procedures with 0.25% Trypsin-EDTA (Gibco) and plated either in BioLite 25 cm^2^ vented flasks (Thermo Scientific) to expand and use for further experimentation or on laminin-coated glass coverslips in a 12 well-plate for immunocytochemistry (ICC).

Glass coverslips (12 mm; Fisherbrand) were sterilized with ultraviolet light for six minutes per side and treated with plasma for two minutes using a Harrick Expanded Plasma Cleaner (PDC-002). Then, a 6.7 µg/mL mouse laminin (Corning) solution, diluted in Hank’s Balanced Salt Solution (HBSS; Sigma Aldrich), was applied to the coverslips and incubated at 37°C for 30 minutes. After, the excess solution was removed, and the coverslips were rinsed with sterile water two times. The cells were then seeded on the coverslips, and 1 mL of media was added. The cells were incubated for three days before performing ICC.

SCs were isolated from postnatal day 2 (p2) Sprague Dawley rats based on previously established methods 14. Sciatic nerves were isolated and plated in basic media (Dulbecco’s Modified Eagle Medium [DMEM; Hyclone], 10% (v/v) FBS, 50 U/mL P/S, and 2 mM of L-glutamine [Sigma Aldrich]). SCs were purified from contaminating fibroblasts with an anti-mitotic agent, 10−5 M cytosine arabinoside (ara-CC; Sigma-Aldrich), and complement-mediated cell lysis (anti-CD90/Thy 1.1, Invitrogen). SCs were seeded on laminin-coated coverslips to assess purity, and ICC staining for S100β (Invitrogen: BSM-52506R, Invitrogen: A11035) was performed as detailed below. Spindle morphology and purity >98% was confirmed. Cells were expanded in BioLite 25 cm^2^ vented flasks (Thermo Scientific) in growth media (DMEM, 10% (v/v) FBS, 50 U/mL P/S, 2 mM of L-glutamine, 10 µg/mL bovine pituitary extract [BPE; Gibco], 6.6 *μ*M forskolin [Sigma-Aldrich]) in standard conditions (5% CO2, 37°C). Media was exchanged every 2-3 days.

Central nervous system glia astrocytes were isolated from postnatal day 2 (p2) Sprague Dawley rats based on previously established methods 33,34. Neonatal brains were isolated, and the olfactory bulbs, cerebellum, and meninges were discarded. Brain tissue was suspended in a dissociation media containing DMEM, P/S, and papain (20 U/mL; MP Biomedicals) and manually triturated. After allowing the brain matter to settle for 15 minutes at standard cell culture conditions (5% CO2, 37°C), the liquid supernatant was removed and diluted 1:1 with media (DMEM, 10% FBS, and P/S) to deactivate the papain. The brain matter was resuspended in a second dissociation media containing DMEM, P/S, papain (20 U/mL), and DNase 1 (20 U/mL; Thermo Scientific) and again manually triturated. The liquid supernatant was removed after 15 minutes of incubation and diluted in the same manner as the previous extraction. Nine additional extractions were performed with the first dissociation media (no DNase 1). A total of ten extractions were performed, equal to the total number of brains harvested. The extracted cells were centrifuged at 400 g for five minutes, resuspended in media, and plated on a poly-l-lysine-coated T75 flask (5,000 cells/cm2). After two weeks of culture, the flask was shaken at 200 RPM for four hours to remove microglia on top of the astrocyte monolayer. After removal, the astrocytes were cultured and expanded. ICC staining for GFAP (Invitrogen: PA1-10004, Invitrogen: A11035) was performed to confirm a purity of >98%.

Neuronal explants, dorsal root ganglia (DRG), were isolated from postnatal (p2) Sprague Dawley Rats via previously reported methods 35,36. Whole spines were removed, and connective tissue was trimmed off. Spines were split medially with scissors, and the spinal cord was removed. DRG explants were lifted from the spinal column with forceps and trimmed excess connective tissue with a 15-blade scalpel. DRG were stored in 15 mL tubes filled with Hibernate-A Medium (Gibco) and wrapped with parafilm at 4°C for no more than one week before use.

### Immunocytochemistry

For ICC, all samples were fixed with 4% paraformaldehyde (PFA; Sigma-Aldrich) for 20 minutes and permeabilized with 0.1% (v/v) Triton-X (Sigma-Aldrich) diluted in HBSS for ten minutes at room temperature. Non-specific binding was blocked with freshly prepared 2.5% (v/v) goat serum (Sigma-Aldrich) diluted in HBSS for two hours at room temperature. Primary antibodies and secondary antibody dilutions are listed in Table 1. Primary antibodies were diluted in the 2.5% goat serum solution and incubated at room temperature for one hour, followed by a triplicate rinse of HBSS. Following this step, secondary antibodies were diluted in the 2.5% goat serum solution and incubated at room temperature for one hour. Before imaging, the samples were washed three times with HBSS and mounted on a glass slide with ProLong™ Gold Antifade Mountant with DAPI (Invitrogen). Images were captured with an inverted light microscope (Zeiss Axio Observer).

**Table 1.** Displays Antibodies used for Immunocytochemistry.

| Name | Classification | Dilution | Host Species | Brand | Product # |
| --- | --- | --- | --- | --- | --- |
| **CD105** | M e s e n c h y m a l<br>Marker | 1:250 | Rabbit | BIOSS | BS-0579R |
| **S100β** | Glia Marker | 1:300 | Rabbit | BIOSS | BSM-52506R |
| **Nestin** | Stemness Marker | 1:500 | Mouse | Abcam | Ab254048 |
| **Beta III Tubulin** | Neuronal Marker | 1:300 | Mouse | Sigma | T8660 |
| **GFAP** | Neuronal Marker | 1:250 | Rabbit | Invitrogen | PA1-10004 |
| **Neurofilament-**<br>**Heavy** | Neurite Outgrowth | 1:300 | Mouse | Invitrogen | MA1-2012 |
| **Myelin Basic**<br>**Protein** | Glia Functionality | 1:300 | Rabbit | Invitrogen | MA5-42370 |
| **Alexa Fluor™ 647** | Secondary | 1:1000 | Goat Anti-Mouse | Invitrogen | A32728 |
| **Alexa Fluor™ 546** | Secondary | 1:1000 | Goat Anti-Rabbit | Invitrogen | A11035 |
| **Alexa Fluor™**<br>**488** | Secondary | 1:1000 | Goat Anti-Rabbit | Invitrogen | A27034 |
| **Phalloidin 546** | Cytoskeleton | 1:1000 | - | Invitrogen | A22283 |
| **DAPI** | Nuclei | 1:500 | - | Invitrogen | D1306 |

### Characterizing OM-MSCs with Flow Cytometry

Passage 5 OM-MSC cell populations were characterized using a Beckman Coulter CytoFLEX. A flow buffer solution was prepared in 50 mL of HBSS with 200 µL EDTA (Corning), 50 mg sodium azide (Sigma-Aldrich), and 50 mg of saponin (Sigma-Aldrich). This flow buffer was kept at 4°C for up to four weeks. Cells were passaged with 0.25% Trypsin-EDTA (Gibco) and pelleted. The cell pellet was resuspended in a solution containing a fixable viability stain (Invitrogen; 1 µL per mL of HBSS) and incubated at 4°C for 30 minutes in the dark. The cells were centrifuged at 350 g for five minutes, the supernatant was removed, and the cells were rinsed by resuspension in HBSS to remove excess stain. After centrifuging again to remove the HBSS, the flow buffer supplemented with 5% (v/v) mouse serum (Invitrogen) was added to the rinsed cell pellet and mixed well by gentle pipetting. The cells were incubated in this solution for 10 minutes at 4°C to limit non-specific binding and to permeabilize the cells. Conjugated antibodies (Table 2) were added to the cell solution and mixed well with the suspended cell solution with a pipette. The cells were incubated at 4°C for 30 minutes and then centrifuged at 350 g for five minutes. The supernatant was removed, and the cells were resuspended in a flow buffer. This centrifugation and resuspension were repeated. The cell solution was then run through the flow cytometer. Compensation was established with single fluorescence and unstained controls. Fluorescence minus one (FMO) controls were utilized to determine gating parameters. Analysis was performed with Flowjo™ Software.

**Table 2.** Displays Antibodies/Markers for Flow Cytometry.

| Excitation | E Filter<sup>m</sup> | Marker | Color | Host/Target | Isotype | Clone | Brand | Catalog # | Dilution Factor |
| --- | --- | --- | --- | --- | --- | --- | --- | --- | --- |
| 488 | 585/42 | CD44 | PE | Mouse α-Rat | IgG1 | OX-50 | Bio-Rad | MCA643PE | 1:10 |
| 488 | 525/40 | Nestin | Alexa 488 | Mouse α-Rat | IgG1 | Rat-401 | Abcam | ab197495 | 1:100 |
| 488 | 695/40 | CD90 | PerCP | Mouse α-rat | IgG1 | M R C OX-7 | Abcam | ab216354 | 1:100 |
| 638 | 660/10 | CD45 | Alexa Fluor 647 | Mouse α-Rat | IgG1 | OX-1 | Bio-Rad | MCA43A647 | 1:10 |
| 638 | 780/60 | eFluor 780 Viability | eFluor 780 | All Species | - | - | Invitrogen | 65-0865-14 | 1:1000 |

### Differentiation of OM-MSCs towards SC-like cells

OM-MSCs were differentiated with two separate media compositions: an SC-conditioned media (SCCM) and a growth factor media (GF). OM-MSCs were plated at 5,000 cells/cm^2^ density for both differentiation protocols, kept at standard culture conditions (5% CO2, 37°C) for 21 days, and had complete media exchanges every 2-3 days (Figure 3A). The first protocol utilizes SCCM or media taken directly from SC cultures after incubation for 2-3 days. The second protocol aims to mimic the SCCM by supplementing SC media (DMEM, 10% (v/v) FBS, 50 U/mL P/S, 2 mM of L-glutamine, 10 µg/mL BPE, and 6.6 *μ*M forskolin) with additional forskolin (final concentration of 16.5 μM in media) to increase cyclic adenosine monophosphate (cAMP) levels and three well-known growth factors secreted natively by SCs: 2.5 ng/mL nerve growth factor (NGF; Gibco #13257-019), 2.5 ng/mL glial cell line-derived neurotrophic factor (GDNF; Gibco #PHC7045), 2.5 ng/mL brain-derived neurotrophic factor (BDNF; Gibco #450-02)37,38. Other media formulations were attempted to optimize the differentiation process, such as a previously established differentiation protocol for BM-MSCs to SCs and varying concentrations of forskolin^39^ (Supplementary Fig. 4). ICC was performed with S100β and Nestin primary antibodies to evaluate changes in expression and cell morphology.

To quantify differences in cell shape, CellProfiler™ software was used 40. Ten immunofluorescent image sets (composed of individual channel images of the exact location) were imported in batches per condition to CellProfiler™ and processed through a custom-designed pipeline (Supplemental Figure 5). The Otsu thresholding method identified the cell nuclei from the DAPI channel. In combination with the identified nuclei and a secondary channel (S100β), the cell body was identified through a propagation method and Otsu thresholding. The “MeasureObjectSizeShape” function characterized each cell body, and results were outputted into an Excel file.

### Gene Expression Evaluation with qPCR

Gene expression was evaluated using RT-qPCR, comparing RNA extracted from DRG, astrocytes, SCs, OM-MSCs, and dOM-MSCs (SCCM and GF media). RNA was collected and isolated from cell pellets with an RNeasy® Mini Kit (Qiagen: 74106) following the manufacturer’s instructions. In place of the gDNA eliminator column, the RNase-Free DNase Set (Qiagen: 79254) was used to ensure sample purity. Ethanolytic precipitation was performed after isolating the RNA to remove excess salts and contaminants. Two volumes of 100% ethanol and 0.1 volumes of 3M sodium acetate (Invitrogen: AM9740) were added to each sample. After vortexing, the samples were kept at -80°C. Before translating the RNA to complementary DNA (cDNA), the RNA was pelleted by centrifugation for 20 minutes (>8000 x g, 4°C). Excess liquid was removed, and the pellet was washed with 100% ethanol and centrifuged again for 10 minutes (>8000 x g, 4°C). The pellet was washed and spun down one more time. After removing all liquid from the pellet, the RNA was allowed to air dry on ice for 10-15 minutes. RNASE-DNASE-free water (12 μL) was added to the pellet to resuspend the RNA. Yield and purity were quantified with a Nanodrop 1 (Thermo Scientific, Supplemental Table 1). Samples with an A260/A280 value between 1.95 and 2.1 were translated to cDNA. The Quantitect Reverse Transcription Kit (Qiagen: 205311) was used according to the manufacturer’s instructions to translate RNA to cDNA, and the cDNA was stored at -20°C.

Primers for genes of interest were designed by inputting the National Center for Biotechnology Information (NCBI) reference sequence into Primer-BLAST41 (Table 3, Supplementary Table 2). Preference was given to primers with a PCR product of 70-200 base pairs (bp) and a melting temperature between 58-62°C. Lyophilized, salt-free primer pairs were then ordered through Eurofin. These primers were then resuspended in Tris-EDTA (TE; Fisher Scientific) to a stock concentration of 10 μM. A standard curve was performed to validate primers with a 10x serial dilution of cDNA starting at a concentration of 1 ng/μL. Optimal primer concentrations were determined based on standard curve efficiency (Supplementary Table 3).

**Table 3:**
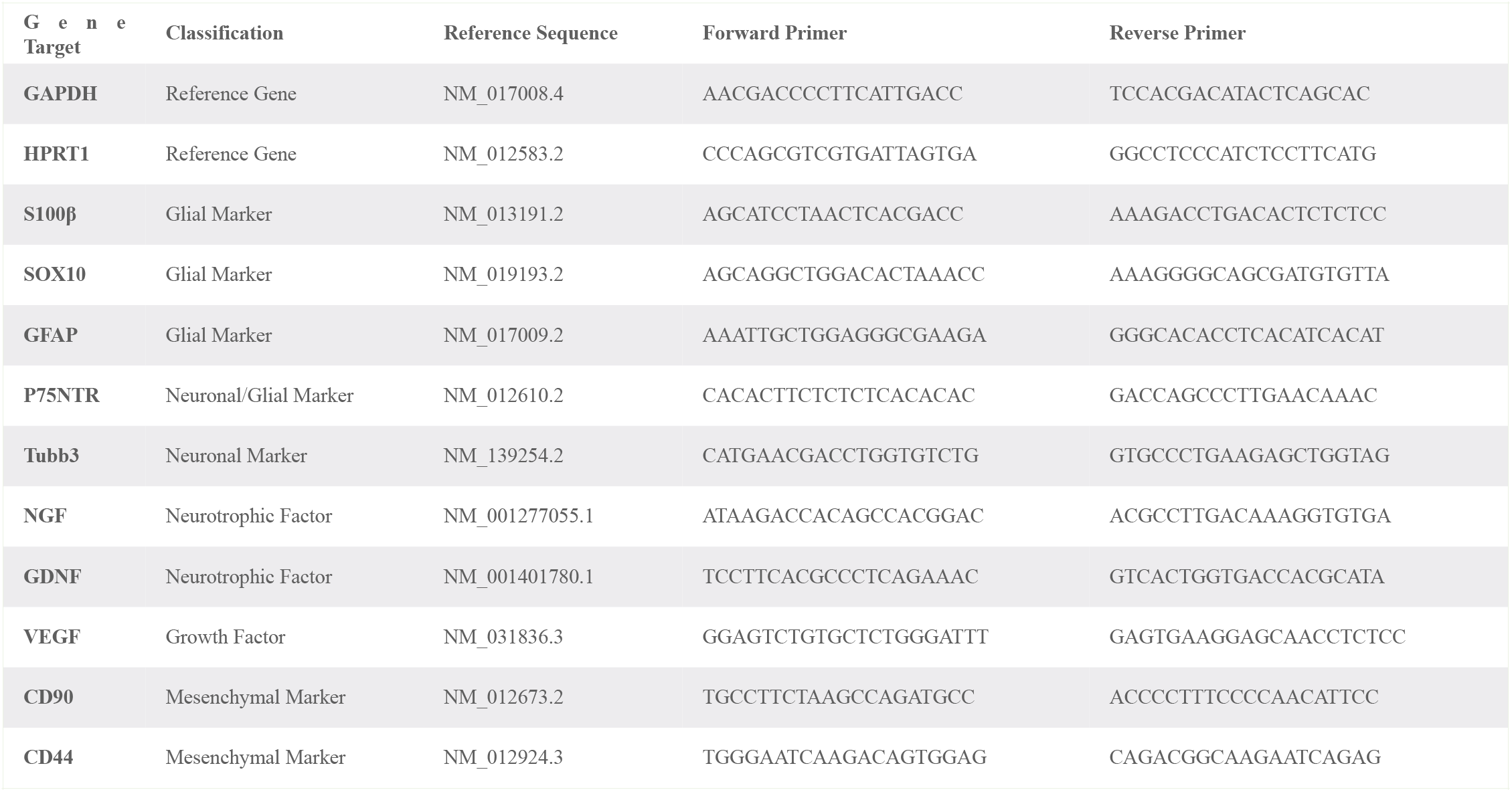
Displays Gene Targets and Primer Sequences.

The expected PCR product size was calculated using AmplifX 42, and the PCR product produced was verified with gel electrophoresis (Supplemental Table 3, Supplemental Figure 6A-B). Briefly, 2% (w/v) agarose gel was made by heating and dissolving agarose (Invitrogen) in 1x Tris Acetate EDTA (TAE; Thermo Scientific). SYBR® Safe DNA gel stain (10,000x: Invitrogen) was added to the solution to a final concentration of 1x, and the solution was then poured into a gel mold (Invitrogen), and a well comb was added. After solidification, the well comb was removed, and the gel was placed in the horizontal gel electrophoresis system. 1x TAE buffer with DNA gel stain was used to submerge the gel. Samples and a 50 bp DNA ladder were loaded into wells with Loading Dye (Invitrogen). The gel was run at 60 V at 0.03 A for two hours with a PowerEase® 300W (Life Technologies). Gels were imaged immediately after the run with a Gel Doc™ XR+ (Bio-Rad). Image Lab 6.1 (Bio-Rad) software was used to calculate the PCR product size.

Eight reference gene primers were designed and validated: beta-2-microglobin (B2M), glyceraldehyde-3-phosphate dehydrogenase (GAPDH), glucuronidase beta (GUSB), hypoxanthine-guanine phosphoribosyl transferase (HPRT1), ornithine decarboxylase antizyme 1 (OAZ1), peptidylprolyl isomerase A (PPIA), ribosomal protein lateral stalk subunit P0 (RPLP0), and ribosomal protein S13 (RSP13) (Supplementary Table 2). A gene study was performed across experimental groups using CFX Maestro software (Bio-Rad) with five validated reference genes (GAPDH, HPRT1, GUSB, RSP13, and B2M) to confirm stability as a reference gene. Samples from the six experimental groups were pooled together to assess the stability. GAPDH and HPRT1 had the highest stability and were chosen as reference genes to use in the experimental quantification plates (Supplementary Fig. 6C). Ten genes of interest were designed, validated, and evaluated: S100 calcium-binding protein B (S100β; glial cell marker), SRY-box transcription factor 10 (SOX10; glial cell marker), glial fibrillary acidic protein (GFAP; glial marker), nerve growth factor receptor (P75NTR; neurotrophin receptor), tubulin beta 3 class III (TUBB3; neural marker), nerve growth factor (NGF; neurotrophic factor), glial cell-derived neurotrophic factor (GDNF; neurotrophic factor), vascular endothelial growth factor (VEGF; growth factor), cluster of differentiation 90 (CD90/Thy-1 cell surface antigen; mesenchymal stem cell marker), and cluster of differentiation 44 (CD44; (mesenchymal stem cell marker) (Supplementary Table 3).

Three 384-well (Thermo Fisher) quantification plates were designed and executed, each containing the two reference genes and experimental genes to minimize intraplate variability. 4 ng of cDNA, 5 μL of SsoAdvanced™ Universal SYBR® Green Supermix, and one set of forward/reverse primer pairs (250-500 nM) were added to each well. All reactions were run in triplicate with four samples per experimental group. After the plate setup, the plate was sealed with Absolute qPCR Plate Seals (Thermo Scientific) and spun down for one minute. PCR was run with a CFX384 thermal cycler (Bio-Rad). 40 cycles of denaturation (98°C, 10 seconds), annealing (60°C, 20 seconds), and extension (72°C, 20 seconds) were performed. A melt curve was also generated with a temperature gradient of 65-95°C at 0.1°C per second. Gene expression was normalized to the two reference genes, GAPDH and HPRT1, and changes in relative expression (log2fold) were quantified using the ΔΔCq method43.

### Co-culture with DRG to Assess the Functionality of Differentiated Cells

Dissociated DRG were cultured with SCs, OM-MSCs, dOM-MSCs (SCCM), and dOM-MSCs (GF) to assess the ability of our differentiated cells to myelinate neurons. The procedure is detailed in the timeline in Figure 6A and is based on previous protocols 44,45. DRG explants were dissociated in 1 mg/mL collagenase (Gibco), 1 mg/mL dispase (Gibco), 15 U/mL papain (Sigma-Alrich) in HBSS for 45 minutes and then triturated 20-40 times with a glass fire-polished pipette. The solution was then diluted at a 1:1 ratio with Neurobasal Media-A (Gibco) to stop the enzymatic activity and centrifugated at 400 g for five minutes. The supernatant was removed, and the dissociated cells were resuspended in Neurobasal-A media, supplemented with 2 mM L-glutamine, 50 U/mL P/S, 1x B-27 (Gibco), 25 ng/mL NGF, and 50 µg/mL of ascorbic acid (Sigma-Aldrich) and plated on laminin-coated coverslips in 24 well plates at a density of 40,000 cells per well. After 12 hr, dissociated DRG were treated with 10−5 M ara-C to reduce the number of native support cells (SCs and fibroblasts). After 24 hr, the ara-C was removed with a full media exchange. SCs, OM-MSCs, and dOM-MSCs (SCCM), or dOM-MSCs (GF), were added to the DRG cultures at a density of 50,000 cells per well. The experiment was maintained in standard conditions (37°C, 5% CO2), and a half media exchange was performed every 3-4 days. The DRG were cultured for 21 days and then fixed and stained as detailed above with mouse anti-neurofilament (NF-H; Invitrogen: MA1-2012) and rabbit anti-myelin basic protein (MBP; Invitrogen: MA5-42370) primary antibodies. Image analysis was performed using a custom-designed CellProfiler™ pipeline, detailed in Supplementary Fig. 7, and the percent pixel overlap of MBP on NF-H was quantified using the “MeasureImageOverlap” function^40^. The Manders coefficient was measured using the “MeasureColocalization” function. Three independent trials were conducted.

### Statistical Analysis

Statistical analysis was performed with Prism GraphPad. The normality of each data set was assessed with the Shapiro-Wilk Test. For parametric (gene expression) data, significance was evaluated with one-way ANOVA and Tukey’s multiple comparison test. For nonparametric data (CellProfiler™ data analysis), a Kruskal Wallis test with a post hoc Dunn’s multiple comparison test was performed. Error bars on graphs represent the standard deviation (SD). *, **, ***, and **** represent a p-value of less than 0.05, 0.01, 0.001, and 0.0001, respectively.

## Data availability statement

The data supporting this study’s findings are available upon reasonable request from the authors.

## Acknowledgments

The authors would like to thank Dr. Rebecca Willits and Dr. McKay Cavanaugh for their assistance and use of their flow cytometer and Dr. David Diaz for his help isolating astrocytes and Schwann cells. Additionally, we would like to thank the Center for Research Innovation (CRI) and Spark Fund program at Northeastern University for all their support.

## Author Contributions

K.N. and R.K. conceived the project. K.N. performed all experimental work analysis and wrote the manuscript. R.K. and A.K. provided analysis support. A.K. and R.K. provided intellectual input and advice. All authors edited and provided feedback on the manuscript. R.K. supervised the work.

## Additional Information

**Supplementary** Information accompanies this paper attached below.

## Supplementary Information

**Supplemental Figure 1.**
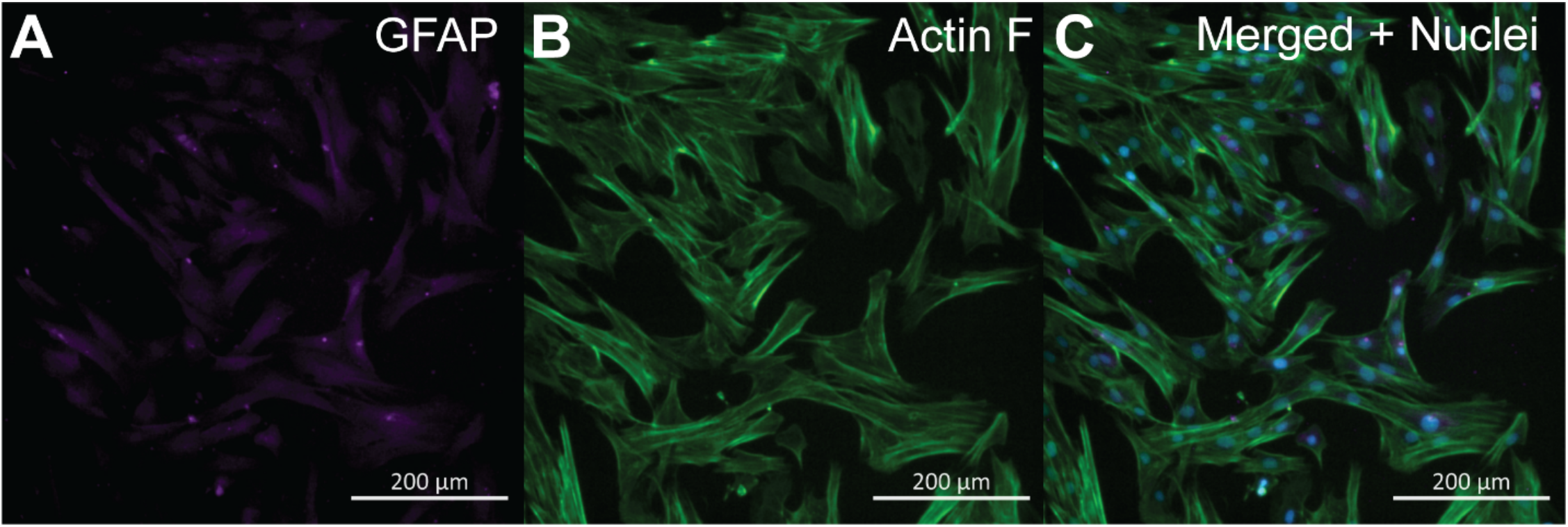
Characteristic immunocytochemistry of OM-MSCs. (A) Purple fluorescence denotes GFAP, (B) Green fluorescence denotes actin filaments, and (C) Displays both GFAP and actin filament expression merged with cell nuclei (shown in blue, counterstained with DAPI). Scale Bar = 200 µm.

**Supplemental Figure 2.**
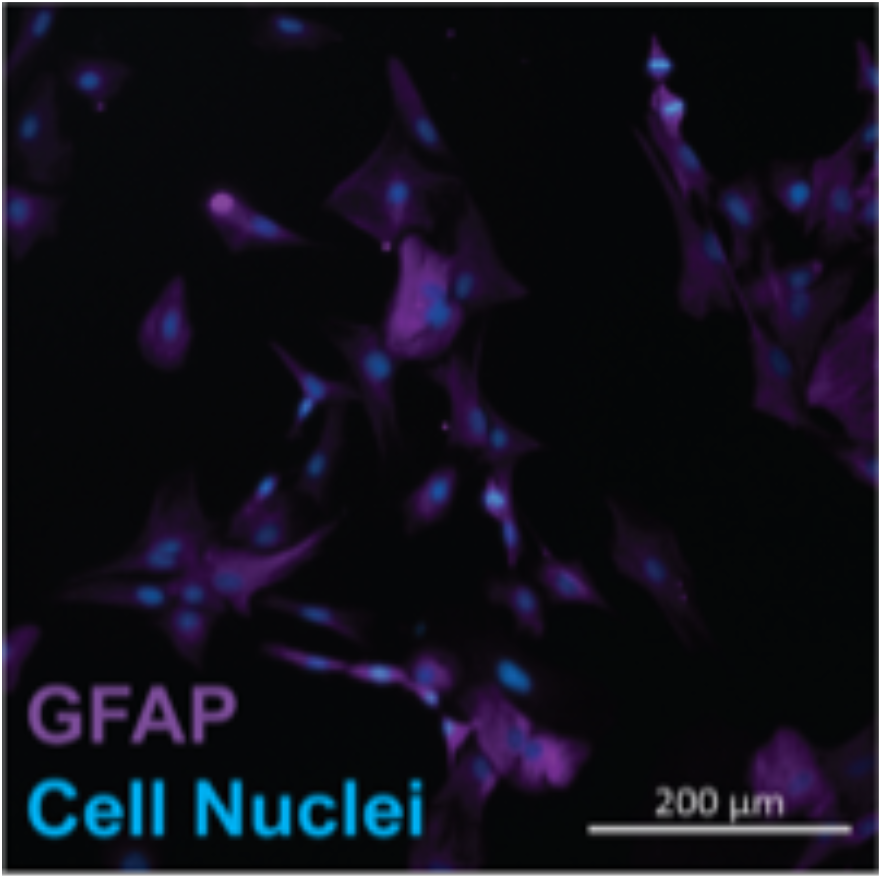
Characteristic immunocytochemistry of isolated astrocytes. Purple fluorescence denotes GFAP expression, and blue fluorescence denotes cell nuclei (counterstained with DAPI). Scale Bar = 200 µm.

**Supplemental Figure 3.**
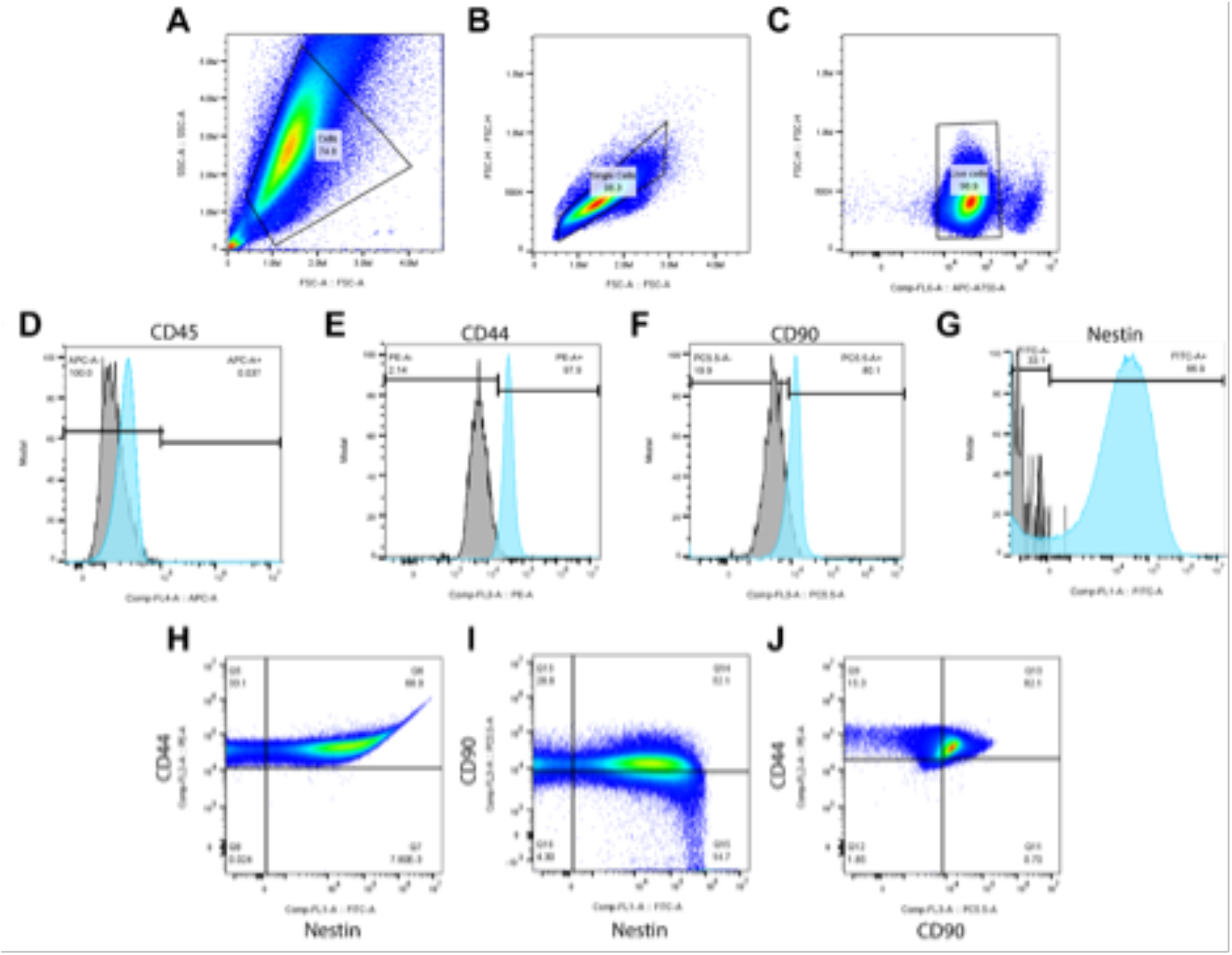
Characteristic flow cytometry of OM-MSCs isolated from adult male Sprague-Dawley rats. (A-C) Display gating strategy employed before analysis. The selection of (A) cell population and removal of debris, (B) single cells, and then (C) live cell population is shown. (D-G) Display the distribution of positive and negative cells for the four markers of interest: (D) CD45, (E) CD44, (F) CD90, and (G) nestin. The gray histogram represents the negative control, and the blue histogram represents the cells of interest. (H-I) Display the co-localization of (H) CD44 and nestin expression, (I) CD90 and nestin expression, and (J) CD44 and CD90 expression.

**Supplemental Figure 4.**
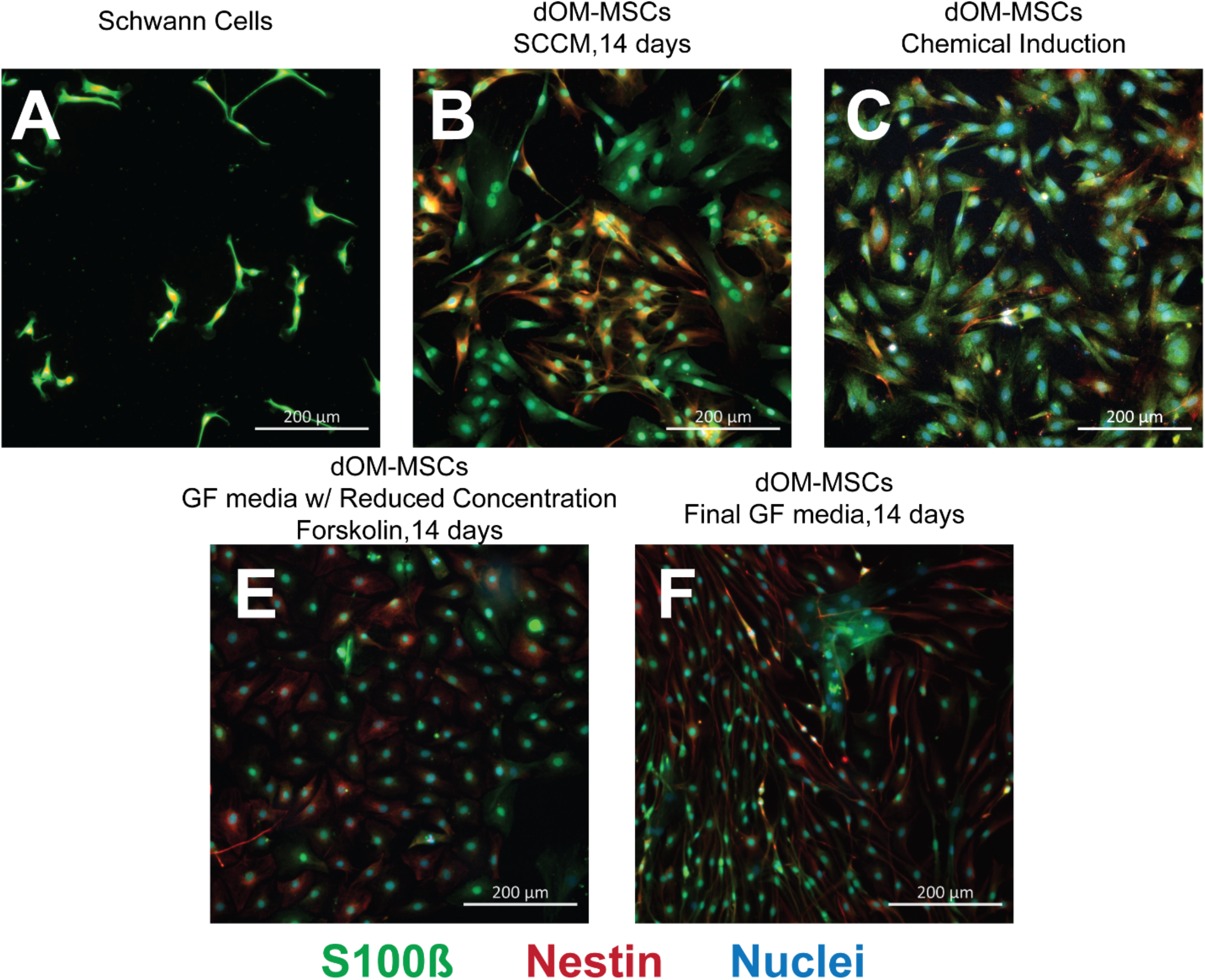
Morphology and expression of S100β and Nestin for (A) Schwann cells, (B) OM-MSCs after culture in SCCM for 14 days, (C) OM-MSCs after a chemical induction protocol, (E) OM-MSCs after culture in GF media with reduced forskolin for 14 days, and (F) OM-MSCs after culture in GF media for 14 days. Green fluorescence indicates S100β, red fluorescence indicates Nestin, and blue fluorescence indicates the cell nuclei after counterstaining with DAPI. Scale bar = 200 µm.

**Supplemental Figure 5.**
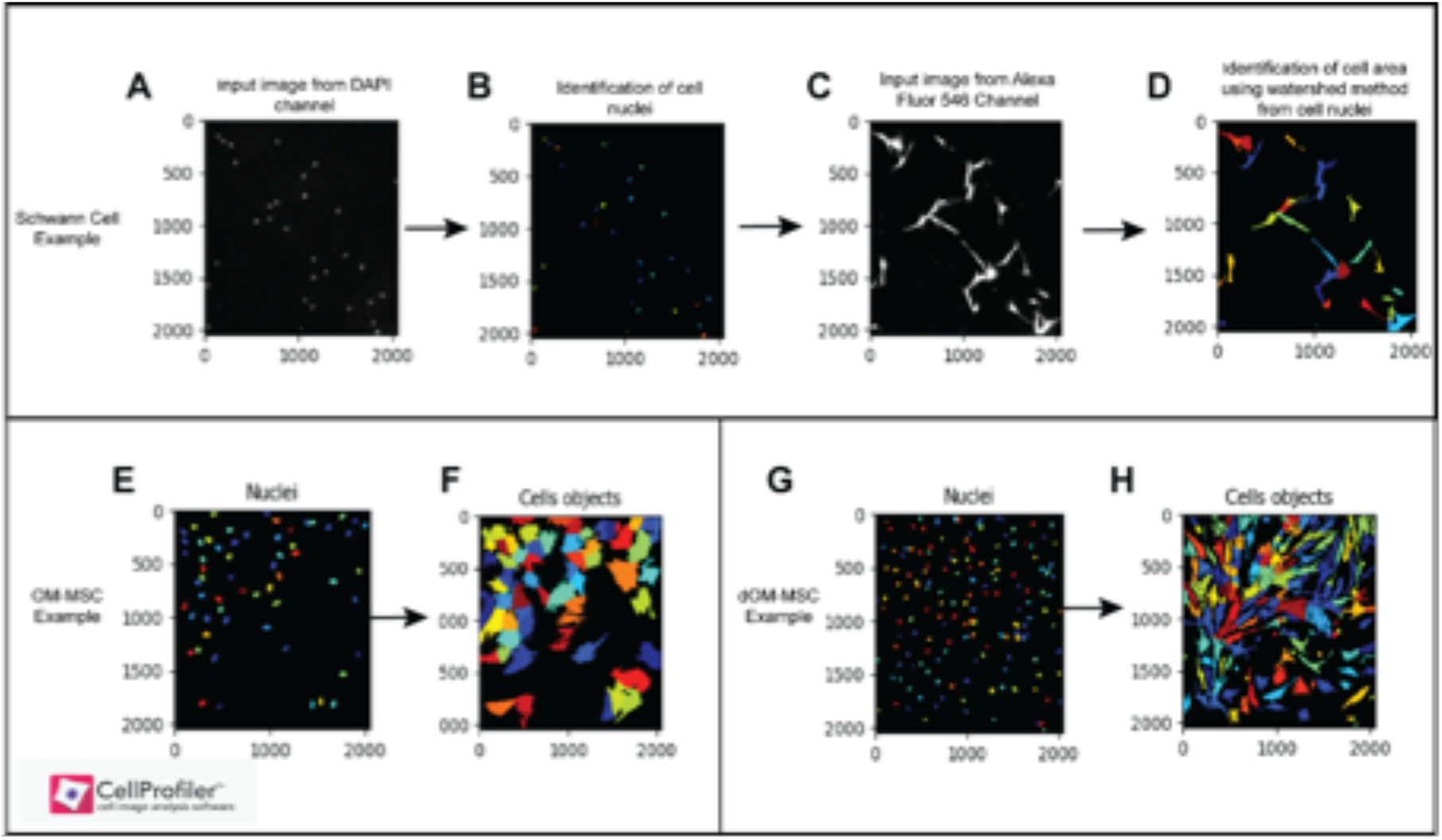
Cell Profiler™ Pipeline examples for (A-D) SCs, (E-F) OM-MSCs, and (G-H) dOM-MSCs. The cell nuclei are identified from the DAPI channel and used to identify and analyze the cell body.

**Supplemental Table 1.** mRNA Yield and Purity for RT-qPCR.

| Sample Name | RNA Yield<br>(ng) | A260/A280 |
| --- | --- | --- |
| SC 1 | 11,126 | 2.1 |
| SC 2 | 15,230 | 2.1 |
| SC 3 | 10,199 | 2.06 |
| SC 4 | 15,127 | 2.07 |
| DRG 1 | 5,915 | 2.05 |
| DRG 2 | 5,300 | 2.03 |
| DRG 3 | 9,400 | 2.08 |
| DRG 4 | 2,652 | 1.98 |
| Astrocyte 1 | 2,929 | 1.95 |
| Astrocyte 2 | 10,600 | 2.04 |
| Astrocyte 3 | 9,900 | 2.02 |
| Astrocyte 4 | 19,153 | 2.09 |
| OM-MSC 1 | 13,500 | 2.12 |
| OM-MSC 2 | 6,100 | 2.04 |
| OM-MSC 3 | 5,400 | 1.98 |
| OM-MSC 4 | 7,908 | 1.97 |
| dOM-MSC 1 (SCCM) | 1,844 | 1.9 |
| dOM-MSC 2 (SCCM) | 1,266 | 2.0 |
| dOM-MSC 3 (SCCM) | 2,974 | 1.98 |
| dOM-MSC 4 (SCCM) | 1,670 | 2.08 |
| dOM-MSC 1 (GF) | 3,380 | 2.01 |
| dOM-MSC 2 (GF) | 3,380 | 1.95 |
| dOM-MSC 3 (GF) | 4,350 | 2.05 |
| dOM-MSC 4 (GF) | 12,000 | 1.98 |

**Supplemental Table 2.**
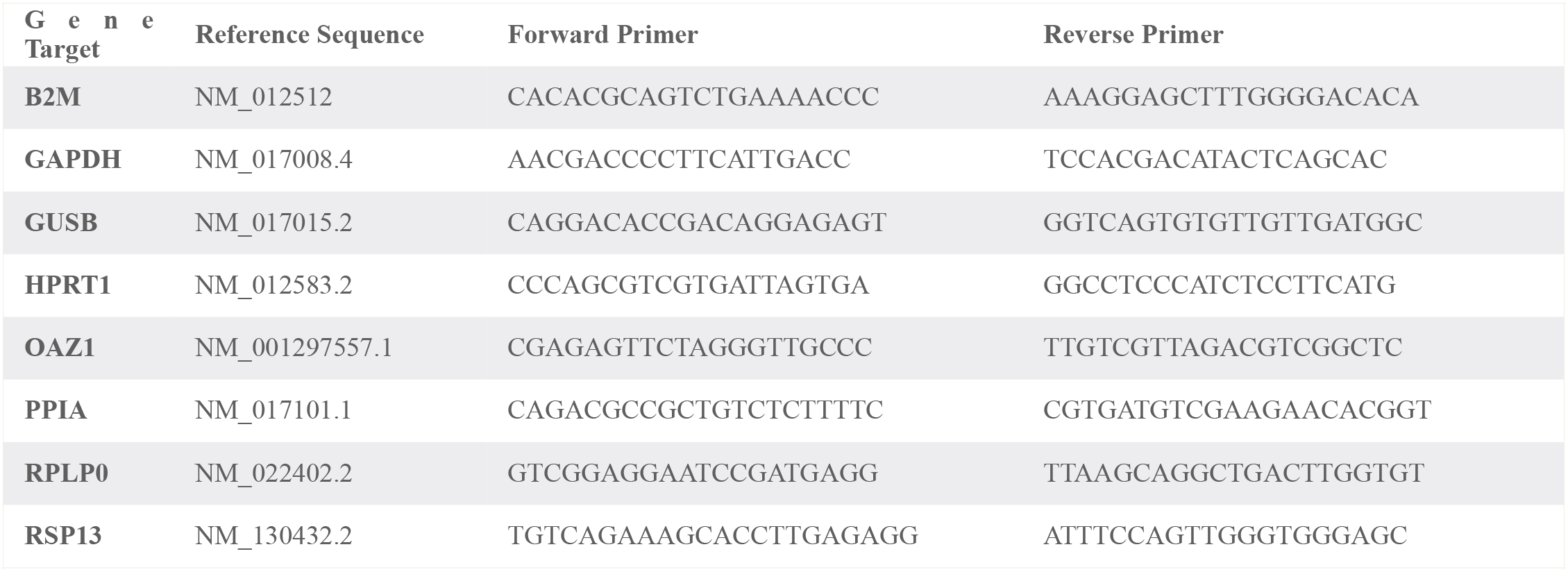
Reference Genes and Primer Sequences.

**Supplemental Table 3.** Validation of Primer Pairs.

| Gene Target | Primer Concentration (nM) | STD Curve Efficiency (%) | R<sup>2</sup> | Expected Band Size (bp) | Estimated Band Size (bp) |
| --- | --- | --- | --- | --- | --- |
| B2M | 500 | 92.5 | 0.995 | 184 | 167 |
| GAPDH | 500 | 96.5 | 0.996 | 191 | 190 |
| GUSB | 250 | 108.7 | 1.000 | 192 | 194 |
| HPRT1 | 500 | 95.8 | 0.999 | 163 | 158 |
| OAZ1 | 500 | 105.6 | 0.997 | 196 | 196 |
| PPIA | 500 | 91.3 | 0.991 | 70 | 67 |
| RPLP0 | 500 | 105.8 | 0.997 | 74 | 73 |
| RSP13 | 500 | 91.3 | 0.998 | 131 | 132 |
| S100β | 500 | 97.1 | 0.990 | 121 | 121 |
| SOX10 | 500 | 92.9 | 0.998 | 116 | 129 |
| GFAP | 500 | 90.3 | 0.984 | 207 | 199 |
| P75NTR | 250 | 104.8 | 0.998 | 251 | 249 |
| Tubb3 | 375 | 96.7 | 1.00 | 239 | 228 |
| NGF | 750 | 90.2 | 0.995 | 198 | 190 |
| GDNF | 500 | 95.5 | 0.991 | 171 | 168 |
| VEGF | 500 | 102.5 | 0.978 | 161 | 132 |
| CD90 | 500 | 100.7 | 0.996 | 279 | 284 |
| CD44 | 500 | 98.1 | 0.991 | 117 | 119 |

**\*\*STD Curve = Standard Curve**

**Supplemental Figure 6.**
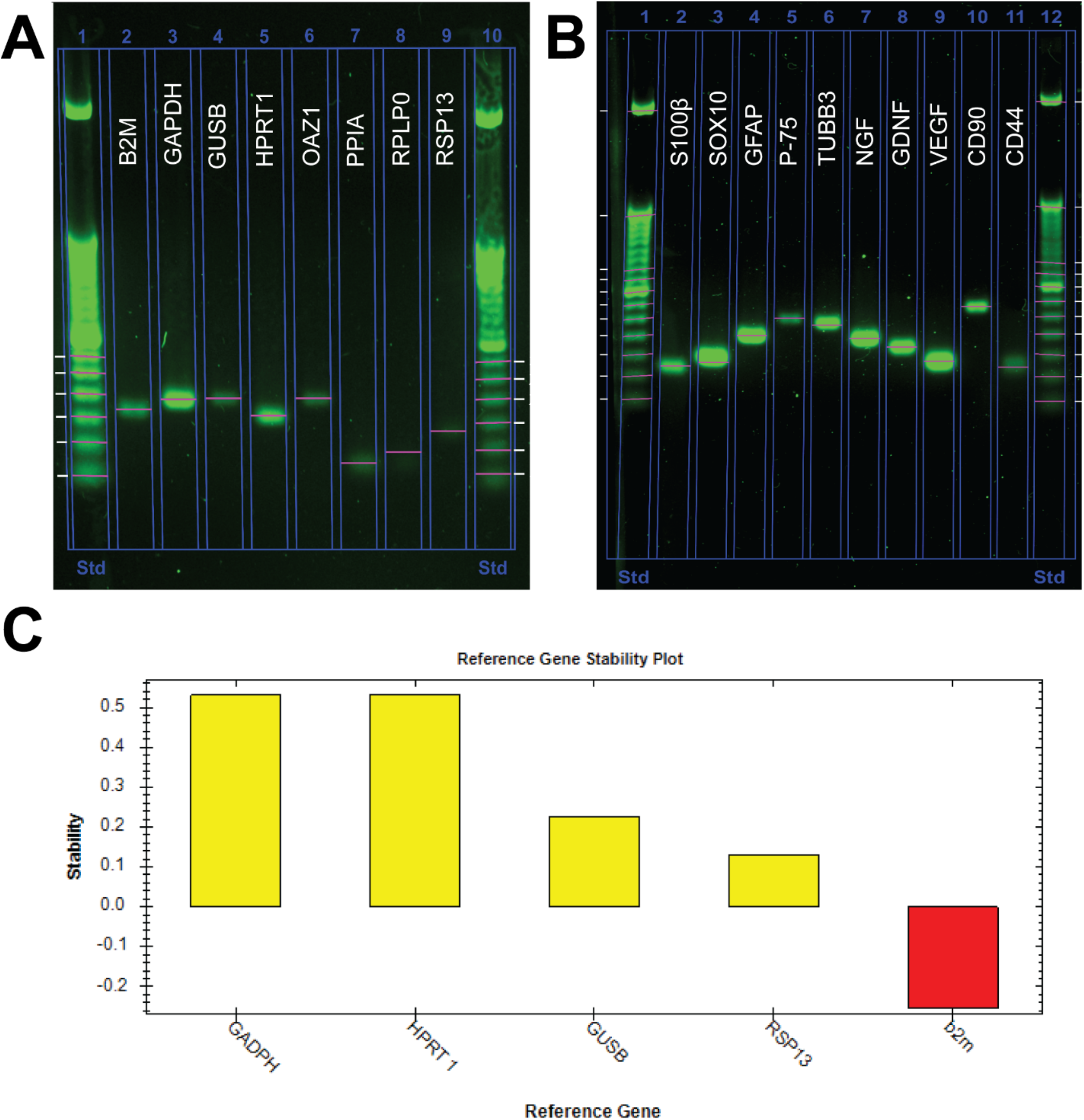
**(A)** Band size verification of PCR products for reference genes with gel electrophoresis. **(B)** Band size verification of PCR products for experimental genes with gel electrophoresis. **(C)** The stability of reference genes were assessed with CFX Maestro software (Bio-Rad).

